# Time-reversal symmetric component of the flagellar beat enhances the translational and rotational velocities of isolated Chlamydomonas flagella

**DOI:** 10.1101/2021.05.01.442280

**Authors:** A. Gholami, R. Ahmad, A. Bae, A. Pumir, E. Bodenschatz

**Affiliations:** Max Planck Institute for Dynamics and Self-Organization, Am Fassberg 17, D-37077 Göttingen, Germany; Department of Biomedical Engineering, University of Rochester, USA; Laboratoire de Physique, Ecole Normale Suprieure de Lyon, Universit Lyon 1 and CNRS, F-69007 Lyon, France; Institute for Nonlinear Dynamics, University of Göttingen, D-37073 Göttingen, Germany; Laboratory of Atomic and Solid-State Physics and Sibley School of Mechanical and Aerospace Engineering, Cornell University, Ithaca, New York 14853, USA

**Keywords:** Chlamydomonas reinhardtii, Flagella wave dynamics, Mode analysis

## Abstract

The beating of cilia and flagella is essential to perform many important biological functions, including generating fluid flows on the cell surface or propulsion of micro-organisms. In this work, we analyze the motion of isolated and demembranated flagella from green algae *Chlamydomonas reinhardtii*, which act as ATP-driven micro-swimmers. The waveform of the *Chlamydomonas* beating flagella has an asymmetric waveform that is known to involve the superposition of a static component, corresponding to a fixed, intrinsic curvature, and a dynamic wave component traveling in the base-to-tip direction at the fundamental beat frequency, plus higher harmonics. Here, we demonstrate that these modes are not sufficient to reproduce the observed flagella waveforms. We find that two extra modes play an essential role to describe the motion: first, a time-symmetric mode, which corresponds to a global oscillation of the axonemal curvature, and second, a secondary tip-to-base wave component at the fundamental frequency that propagates opposite to the dominant base-to-tip wave, albeit with a smaller amplitude. Although the time-symmetric mode cannot, by itself, contribute to propulsion (scallop theorem), it does enhance the translational and rotational velocities of the flagellum by approximately a factor of 2. This mode highlights a long-range coupled on/off activity of force-generating dynein motors and can provide further insight into the underling biology of the ciliary beat.

## 1. Introduction

Cilia and flagella are slender hair-like appendages that protrude from the cell surface and act as a fundamental motility unit by performing periodic whip-like motion to provide driving force for fluid transport or cell locomotion [1, 2]. Typical examples are mucuciliary clearance in mammalian airways to protect the respiratory system from harmful inhaled materials [3, 4], transport of cerebrospinal fluid in the brains of mammals [5], transport of the egg to the uterus [6], cilia-driven flow determining the left-right asymmetry in the embryonic node [7], or propulsion of micro-organisms such as paramecium, spermatozoa, or the unicellular biflagellate alga *Chlamydomonas reinhardtii* (*C. reinhardtii*) [8–12].

The core structure of cilia, known as axoneme, has a highly conserved cylindrical architecture consisting of nine microtubule doublets at the periphery and two microtubule singlets at the center [13–15]. The diameter of axoneme is about 200 nm and doublet microtubules (DMTs) are spaced 30 nm away from each other (see Fig. 1). To actuate a flagellum, dynein molecular motors which are bound periodically in two rows to the DMTs, convert efficiently chemical energy from ATP hydrolysis into mechanical work. One row of the motors are the outer dynein arms (ODAs) which provide power output, and the other row are the inner dynein arms (IDAs) which determine the flagellar beat pattern [16, 17]. For flagella to beat regularly, spatial and temporal mechanical inhibition of dynein molecular motors is required [14]. Multiple ciliary components are involved in this mechanical feedback loop: central pair microtubules (CP), radial spokes (RS) and nexin-dynein regulatory complex (N-DRC). N-DRC is a large, complex structure which interconnects the outer microtubule doublets and maintains their alignment, and is responsible for converting the action of the dynein motors into microtubule bending by limiting microtubule sliding [18, 19]. Radial spokes relay signals from the CP to the dynein arms and guarantee a collective activity of dyneins to form a regular wave pattern [20–23]. Axonemes with defective radial spokes, are either entirely paralyzed or exhibit irregular beat patterns despite having active dyneins [21].

**Figure 1.**
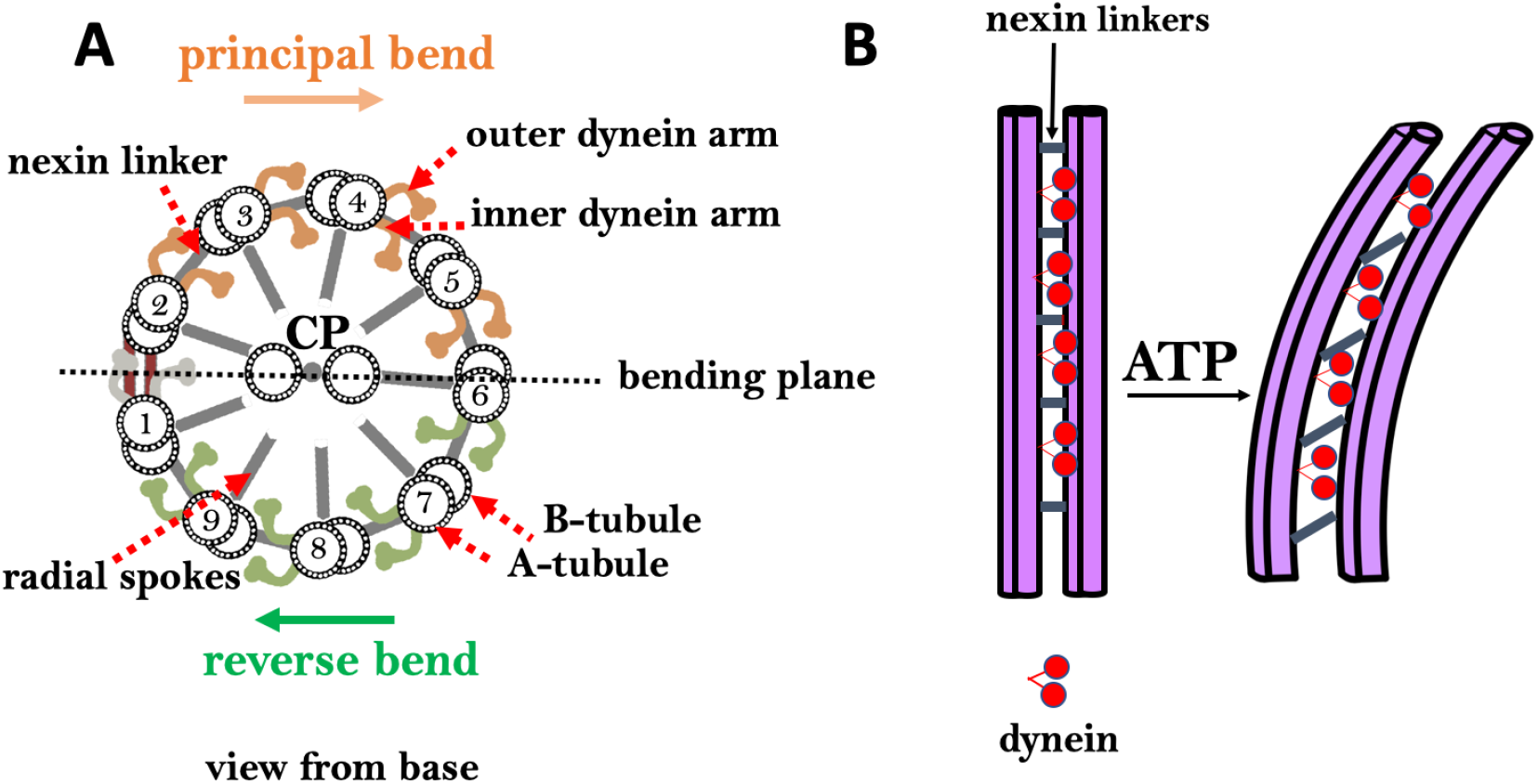
A) 9+2 microtubule structure of ciliary axoneme and the associated flagellar components. B) Dynein motors consume ATP to slide neighboring DMTs relative to each other, but due to the structural constrains which resist sliding, axonemal bending occurs.

The beating pattern of flagella varies among different species. Flagella isolated from green algae *C. reinhardtii* beat by asymmetric curvature waves propagating along the contour length in a base-to-tip direction [24–27]. These asymmetric curvature waves result in a curved swimming trajectory and can be decomposed into multiple mode components [24–26, 28–30]. The magnitude of different components contributing to the flagellar waveform depends on the concentration of calcium ions which are found to be one of the key players in shaping the flagellar beat. Experiments by Bessen et.al. [31, 32] with axonemes isolated from *C.reinhardtii* have shown that high Ca^2+^ concentration triggers a transition from asymmetric to a symmetric waveform. Similar calcium-dependent modification in the wave symmetry was also observed in ATP-reactivated *C. reinhardtii* cells where two flagella are demembranated using detergents [33]. Interestingly, to reverse the direction of wave propagation during a photo-phobic response to an intense light, *C. reinhardtii* cells change the internal level of Ca^2+^ ions to switch from asymmetric forward beating pattern to a symmetric reverse mode of beating [34–37]. Similarly, in ciliated micro-organisms such as *Paramecium* or *trypanosome Crithidia* [38, 39], calcium ions appear to be responsible for avoidance response by altering the wave direction, frequency and bend form. Remarkably, in *trypanosome Crithidia*, curvature waves propagate in the unusual way from the distal tip toward the basal end of the cilium. However, in an avoidance response, a calcium-triggered base-to-tip wave propagation was observed [39].

In isolated and reactivated flagella of *C. reinhardtii*, the following dominant modes in the waveform are observed [24–26]: 1) a semi-circular static mode resulting from averaging the flagellar shapes over a beat cycle, 2) a main dynamic mode describing a base-to-tip wave propagation at the fundamental beat frequency *f*_0_, and 3) a second-harmonic component which describes a base-to-tip wave propagation at frequency 2*f*_0_. The static mode and the second harmonic of the waveform contribute to the axonemal rotational velocity [27, 40, 41] and in the absence of these two components, the axoneme swims in a straight sperm-like trajectory [31, 32]. In this work, we combined high-precision high-speed phase contrast microscopy (1000 fps) with image processing and analytical analysis to study the wave patterns of actively beating axonemes isolated from *C. reinhardtii*. Mode decomposition analysis of our experimental data, demonstrates important contribution of two additional dynamic modes in the flagellar beat: first, a mode corresponding to the global oscillations of the axonemal curvature at the beat frequency *f*_0_ and second, a tip-to-base propagating wave component that also propagates at frequency *f*_0_, but with a reduced amplitude. The mode which describes the global oscillations of the axonemal curvature is time-symmetric and by itself cannot provide propulsion (scallop theorem [42, 43]). However, our analytical analysis confirm that once the other traveling wave components break the time symmetry, these global oscillations contribute in the enhancement of the translational and rotational velocities of the axoneme. Furthermore, to investigate how the constituting wave components change in response to calcium ions, we performed experiments at different calcium concentrations. Our mode decomposition analysis of axonemes reactivated with calcium-supplemented buffer confirm that as we increase the calcium concentration from 0.0001 mM to 1 mM, the static mode drops significantly (~ 85%), triggering a transition from circular swimming path to a straight trajectory. In addition, the main traveling wave component at the fundamental frequency shows a decrease of ~ 40%, while the other components do not exhibit a significant change.

## 2. Results

We isolated flagella from *C. reinhardtii* wild type cells using established protocols [8, 44] and demembranated them using non-ionic detergents. These naked flagella (axonemes), can be reactivated at various ATP concentrations (see Materials and Methods). ATP powers dynein molecular motors that convert chemical energy into mechanical work by sliding adjacent DMTs relative to each other, as illustrated in Fig. 1 [18, 19, 21, 45]. However, structural constrains do not allow DMTs to slide freely. Instead, sliding is converted into rhythmic bending deformations that travel along the contour length of axonemes in the base-to-tip direction. To quantify these curvature waves, we tracked the axonemes over time using the gradient vector flow technique [46, 47] (see Materials and Methods).

Figure 2A-D illustrates planar swimming motion of an exemplary axoneme reactivated at 80 *μ*M ATP (see Video 1). Bending waves initiate at the basal end and travel toward the distal tip of the axoneme at the frequency of *f*_0_ ~18 Hz (Fig. 2C). These traveling periodic curvature waves provide the necessary thrust to propel the axoneme in the surrounding water-like fluid. This thrust force is balanced with the viscous drag exerted by the fluid on the swimmer [48]. The swimming dynamics of axonemes differ from sperm flagellar propulsion primarily in that axonemes are shorter in length (~10 *μ*m in comparison to 50 *μ*m of human spermatozoa) and beat with a nonzero static curvature (Fig. 3A), causing a circular swimming trajectory (Fig. 2B,D). To quantify the static curvature, we translate and rotate the axonemal configurations such that the basal end (*s* = 0) is at the origin of the coordinate system and the tangent vector at *s* = 0 is oriented in the 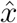 direction (see Fig. 2A). The filament in cyan color in Fig. 3A shows the time-averaged axonemal shape with mean curvature of *κ*_0_ ~ −0.21 *μ*m^−1^. The negative sign of *κ*_0_ indicate a clockwise bend when moving from the basal end at *s* = 0 toward the distal tip at *s* = *L* (Fig. 3A). Base-to-tip bending deformations superimposed on this negative intrinsic curvature cause a counter-clockwise rotation of the axoneme (Fig. 2B).

**Figure 2.**
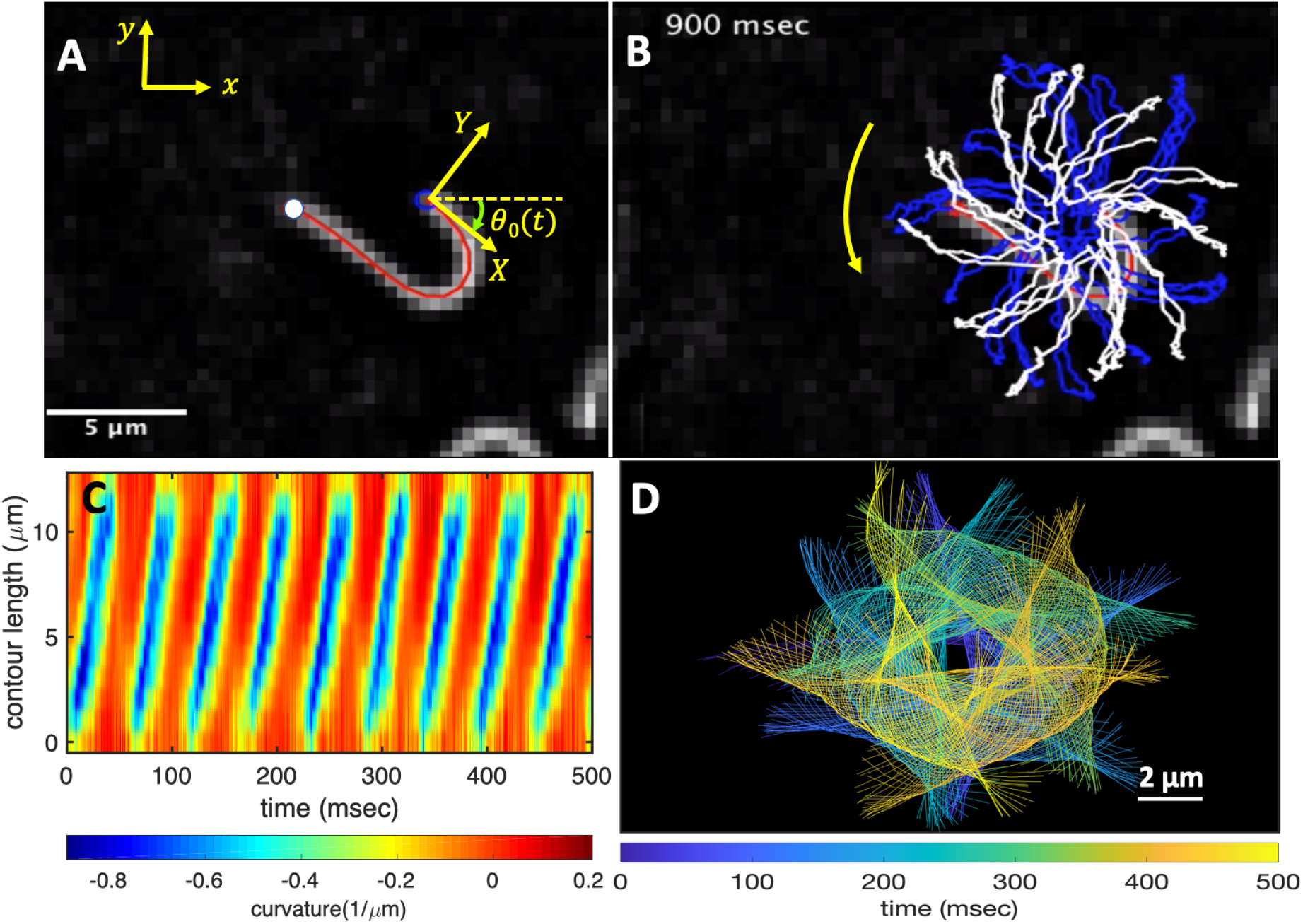
An isolated axoneme beating freely. A) An instantaneous shape. The tangent vector **t**(*s*) at *s* = 0 defines the *X*-direction of the swimmer-fixed frame and the corresponding normal vector **n**(*s*) defines the ***Y***-direction. B) Traces of the basal and the distal ends of the axoneme tracked for 900 msec. C) Curvature waves initiate at the basal end of the axoneme and propagate toward the distal tip at a frequency of 18 Hz. D) The color-coded time projection of the axoneme shows a circular swimming trajectory.

**Figure 3.**
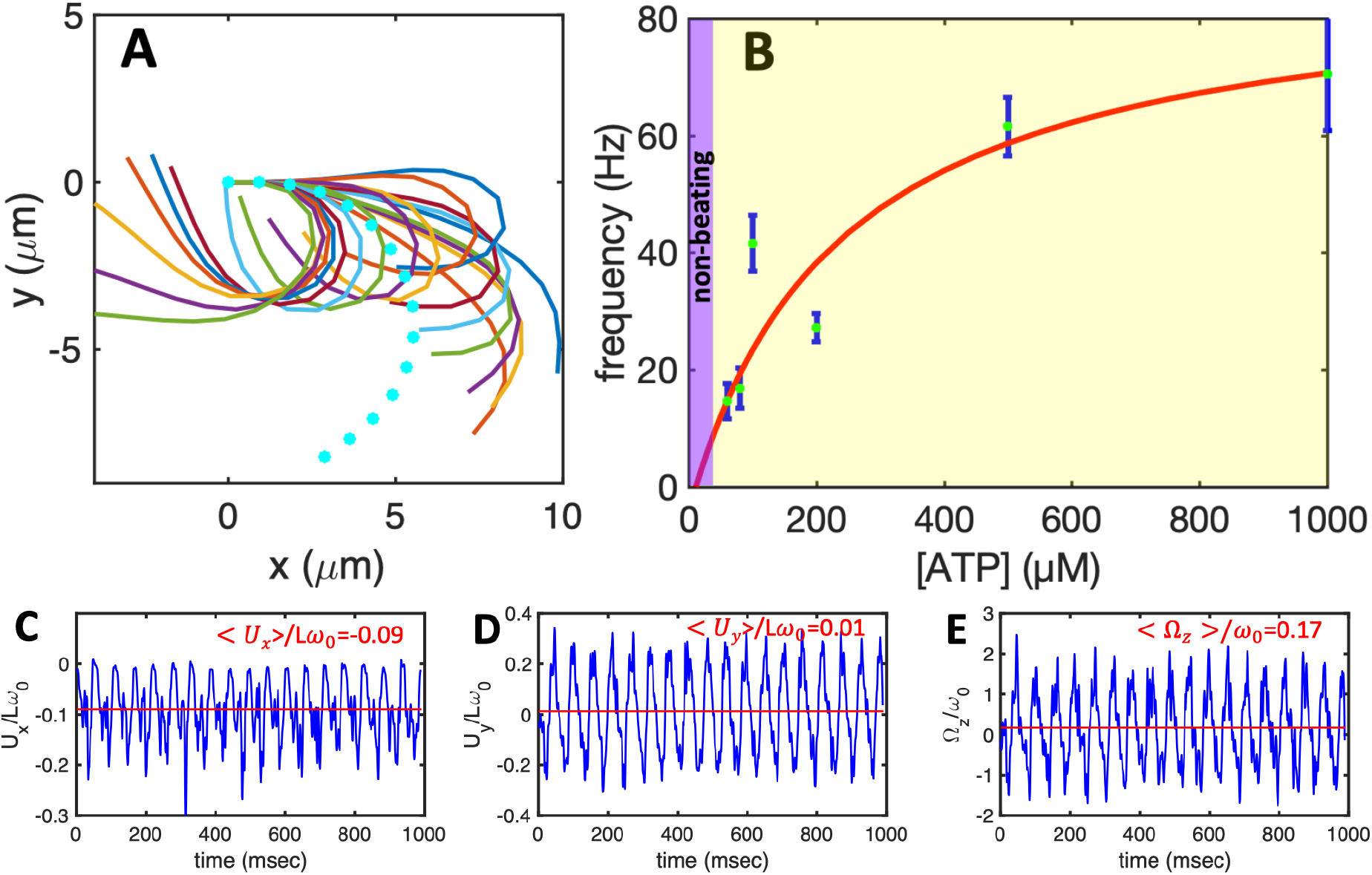
A) The basal end of the tracked axoneme in Fig. 2 is translated to the origin and rotated such that the tangent vector at *s* = 0 is along the *x*-axis. Semi-circular arc in cyan color with mean curvature of *κ*_0_ ~ −0.21 *μ*m^−1^ shows the time-averaged shape of the axoneme. This averaged intrinsic curvature makes the waveform asymmetric. B) Axonemal beating frequency depends on the ATP concentration and is higher at higher ATP concentrations. Axonemes ceased to beat at ATP concentrations below 60 *μ*M ([ATP]_critical_). Red curve shows the least square fit to the Michaelis-Menten relation *f* = *f*_critical_ + *f*_max_([ATP] – [ATP]_critical_)/(*K_m_* + ([ATP] – [ATP]_critical_)). The ftttmg parameters are *f*_max_ = 73.75 Hz and *K_m_*=295.8 *μ*M [26, 49]. C-E) Translational and rotational velocities of the axoneme in Fig. 2, measured in the swimmer-fixed frame.

The beating frequency of axonemes depends on ATP concentration and follows a Michaelis-Menten-type kinetics [26, 49]: it starts with a linear trend at small concentrations of ATP and saturates at higher ATP concentrations around 1 mM (see Fig. 3B). In our experiments, we measured a critical minimum of ATP concentration around 60 *μ*M was required to reactivate axonemes [26, 41]. Reactivated axonemes swim on circular trajectories effectively in 2D (see Ref. [50] for a study on small out-of-plane beating components of isolated axonemes). Active axonemes undergo planar shape deformations over time, but at any instant of time it may be considered as a solid body with translational and rotational velocities *U_x_, U_y_* and Ω_*z*_, which we measure in the swimmer-fixed frame (Fig. 3C-E) [51]. These velocities oscillate in time, reflecting the fundamental beat frequency of the axoneme (18 Hz) and its higher harmonics.

### 2.1. Mode decomposition of the curvature waves

We performed a 2D Fourier analysis of the traveling curvature waves of the reactivated axoneme shown in Fig. 2. For this purpose, we “windowed” the curvature data, meaning that the curvature waves of an arbitrary beat cycle is repeated integer number of times to obtain a fully periodic signal. Exemplarily, we chose the last beat cycle to obtain the periodic “windowed curvature” shown in Fig. S1, and which we use for the following analysis. The corresponding power spectrum demonstrates a dominant peak at zero temporal and spatial frequencies corresponding to the static curvature of the axoneme (see Fig. 4A). This static curvature is denoted by dimensionless number *C*_0_ = *κ*_0_*L*/2*π*, where *L* = 12.35 *μ*m is the contour length of the axoneme. Note that with *κ*_0_ = 0.21 *μ*m^−1^ extracted from Fig. 3A, we expect the mode amplitude *C*_0_ to be around 0.41 (see Fig. 4B). The second dominant peak which occurs at the fundamental beat frequency *f*_0_ (18 Hz) and wavelength λ = *L*, is denoted by *C*_1_ and describes traveling bending deformations which start at the base and propagate toward the distal tip. Surprisingly, the next important peak 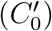 occurs at the beating frequency of 18 Hz but at zero spatial frequency (infinite wavelength). This mode which describes a global oscillations of the axonemal curvature, is invariant under time-reversal and by itself can not generate propulsion (scallop theorem [42, 43]). However, once the time-symmetry is broken through the traveling wave components, it contributes in enhancing the translational and rotational velocities of the axoneme, as we will show in the next section. Moreover, the power spectrum in Fig. 4A also highlights the presence of a less dominant dynamical mode 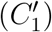 corresponding to a back-propagating wave which starts from the tip and propagates at the frequency of 18 Hz toward the base. Finally, among the other peaks, we also observe a peak at the second harmonic 2*f*_0_ and wavelength *L*. Similar to the static curvature, the second harmonic also contributes to the rotational velocity of the axoneme [27, 40], but the static curvature has the dominant contribution in axonemal rotation (see next section and supplemental information).

**Figure 4.**
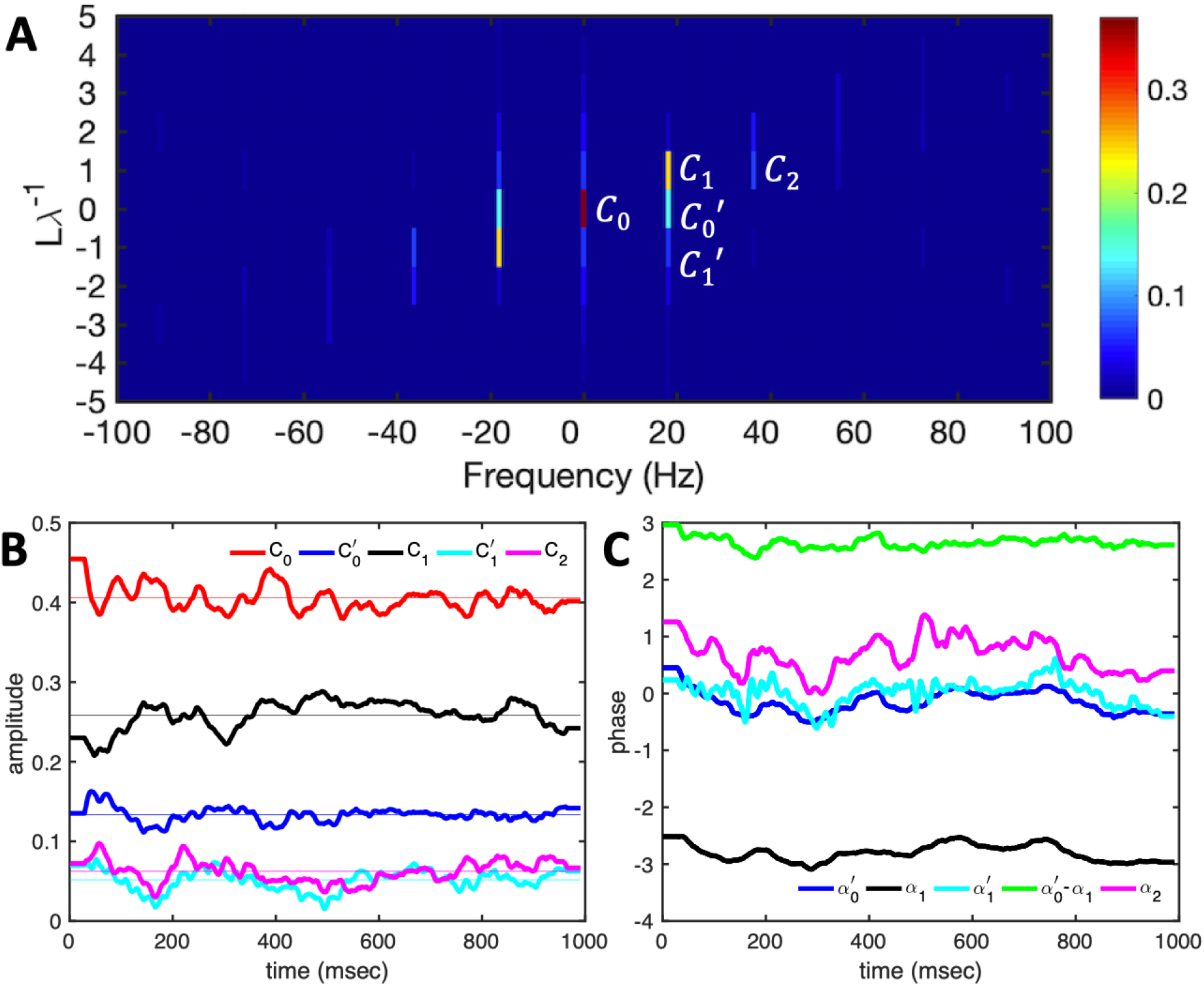
A) Power spectrum of the “windowed” curvature waves shown in Fig. S1. The last beat cycle is used to “window” the curvature. Five dominant modes are highlighted: *C*_0_ = *κ*_0_*L*/2*π* corresponds to the amplitude of the static curvature, 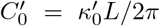 refers to the amplitude of the global oscillations of the curvature, *C*_1_ = *κ*_1_*L*/2*π* is the amplitude of the base-to-tip propagating wave, 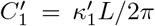 shows the amplitude of the wave propagating in the opposite direction and finally *C*_2_ = *κ*_2_*L*/2*π* is the amplitude of the second harmonic. B-C) The time evolution of the dominant modes and the corresponding phase values: 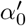 (phase of 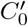), *α*_1_ (phase of *C*_1_), 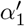 (phase of 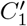) and *α*_2_ (phase of *C*_2_).

The analysis so far, done using only one interval from *t* to *t* + 1/*f*_0_ of the signal, starting at one particular value of *t*, can in fact be generalized by letting the time *t* vary. This allows us to construct periodic ‘windowed” signals, and to determine the corresponding 2D power spectra. The time evolution of the amplitudes of the dominant modes is shown in Fig. 4B. Notably, the mode which corresponds to the global oscillations of the axonemal curvature 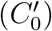, has almost double the amplitude of the second harmonic *C*_2_ and the back-propagating wave component 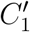. The time-averaged amplitudes of the dominant modes gives 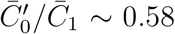, which we will use in our analytical analysis in the next section.

The Fourier analysis also provides important phase information of each contributing mode. Figure 4C shows the time evolution of the phase of the four dynamic modes which are highlighted in Fig. 4A. Remarkably, while the mode describing the global oscillations of the axoneme and the back propagating wave are almost in phase (compare 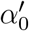 and 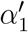), the main dynamic mode and the mode of global oscillations are almost out of phase and show the phase difference of around *π* (compare *α*_1_ and 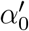).

To describe the axonemal shapes, we used the five dominant modes of the power spectrum shown in Fig. 4B and the corresponding phase information, as described above, setting the other modes to zero. We performed an inverse Fourier transform of the power spectrum to retrieve the curvature waves and reconstruct the shapes. The comparison between experimental shapes and the results obtained by superposition of these five modes, is shown in Fig. 5A-B. The best and worst shape reconstructions correspond to the mean square error of 0.007 and 0.039, respectively, calculated by comparing the experimental with the reconstructed shapes (Fig. 5E-F). To emphasize on the contributing roles of 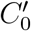 (global oscillations of the curvature), 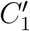 (back-propagating wave) and *C*_2_ (the second harmonic) in shape reconstruction, we also present the results of the shapes reconstructed from *C*_0_, 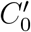, *C*_1_ and 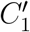 modes without *C*_2_ mode (Fig. 5C and 5G-H), and compared with the shapes reconstructed from *C*_0_, *C*_1_ and *C*_2_ modes, without 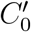 and 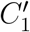 modes (Fig. 5D and 5I-J). It is important to note that while removing the second harmonic does not significantly increase the fitting error, eliminating both 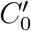 and 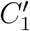 modes increases the error in shape reconstruction more than two times (see also supplemental Videos 2-4). Please note that in the past literature [24, 26] the modes 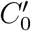 and 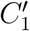 were not considered.

**Figure 5.**
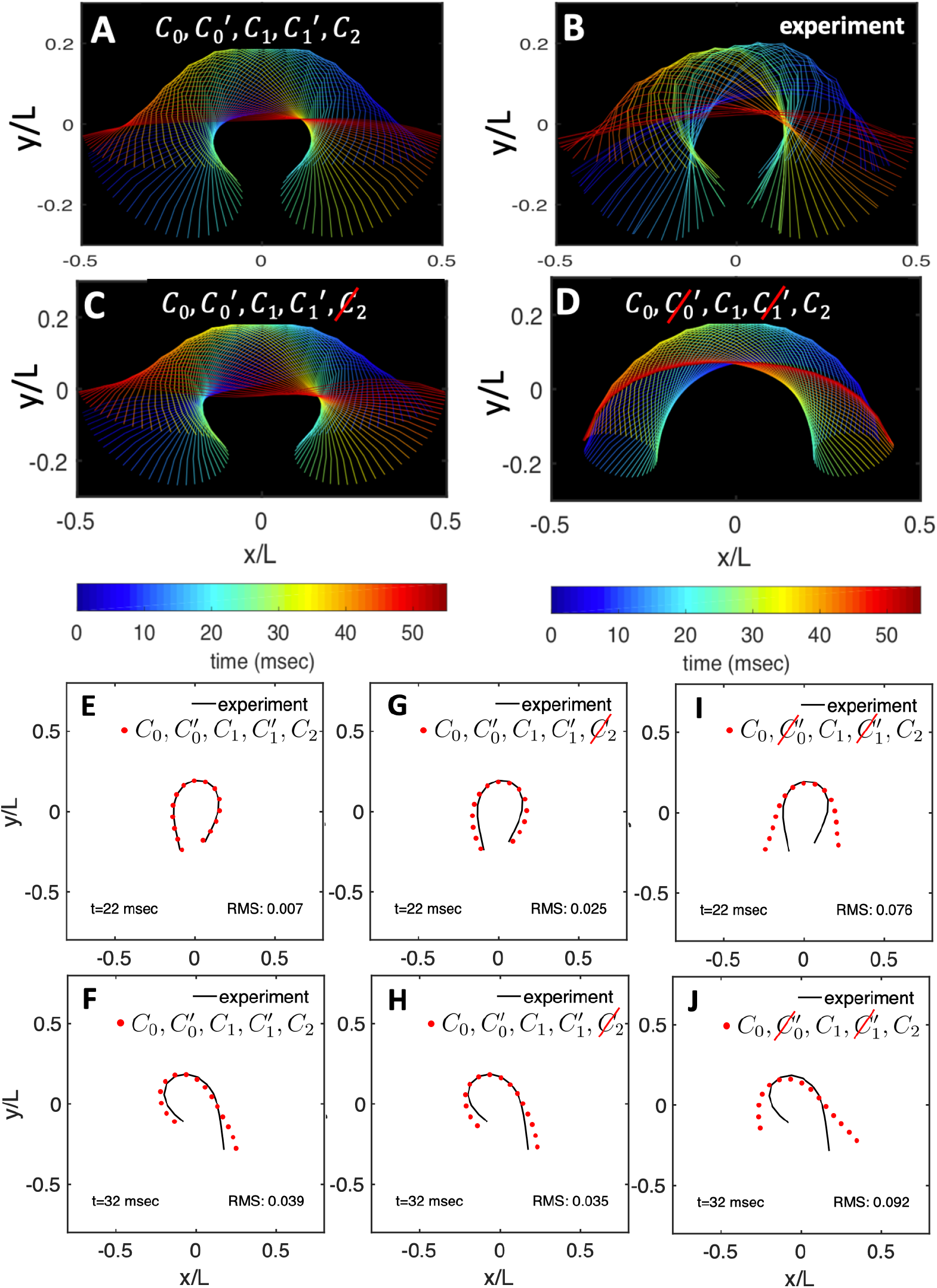
A) Five dominant modes, highlighted in Fig. 4A, are used to reconstruct the experimental shapes shown in panel B. C) The shape reconstruction without the second harmonic *C*_2_ and D) without both back propagating wave 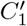 and global oscillation mode 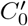. E-F) Exemplary fits of the best and the worst shape reconstructions shown in panel A with the root mean square (RMS) error of 0.007 and 0.039, respectively. G-F) The corresponding time points with 4 modes (no second harmonic) and I-J) with 3 modes (without 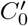 and 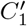) are also presented; see Videos 2-4.

### 2.2. Analytical analysis

To understand the effect of various modes on rotational and translational velocities of a beating axoneme, we first consider the effect of the first four dominant modes, namely the static curvature (*C*_0_), the global oscillations of the curvature 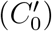, the base-to-tip and tip-to-base propagating waves (*C*_1_ and 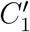):

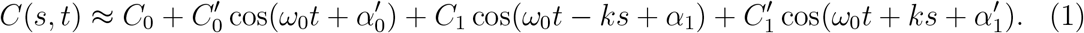

Here *C*(*s,t*) = *κL*/2*π* is the dimensionless curvature at arc-length *s* (0 ≤ *s* ≤ *L*) at time *t, ω*_0_ = 2*πf*_0_, and wave number *k* is defined as *k* = 2*π*/λ ~ 2*π*/*L*, where wavelength is found to be of the order of axonemal length *L*. The effect of the second harmonic is presented in supplemental material and is extensively discussed in Refs. [27, 40]. The amplitudes and phase values are in general time-dependent functions (see Fig. 4B-C), but in our analytical analysis, we assume that these quantities are time-independent.

In the absence of external forces, the total force and torque exerted on a freely swimming axoneme at low Reynolds number regime is zero. Given the prescribed form of curvature waves in Eq. 1, we used resistive force theory (RFT) [52, 53] to calculate propulsive forces and torques from the tangential and normal velocity components of each axonemal segment. By imposing the force-free and torque-free constrains in 2D, we can uniquely determine translational and rotational velocities of the axoneme (see Materials and Methods). We determine analytically the averaged angular and linear velocities of the axoneme in the swimmer-fixed frame up to the first order in *C*_0_ and 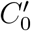 and second order in *C*_1_ and 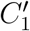 to obtain [27, 40]:

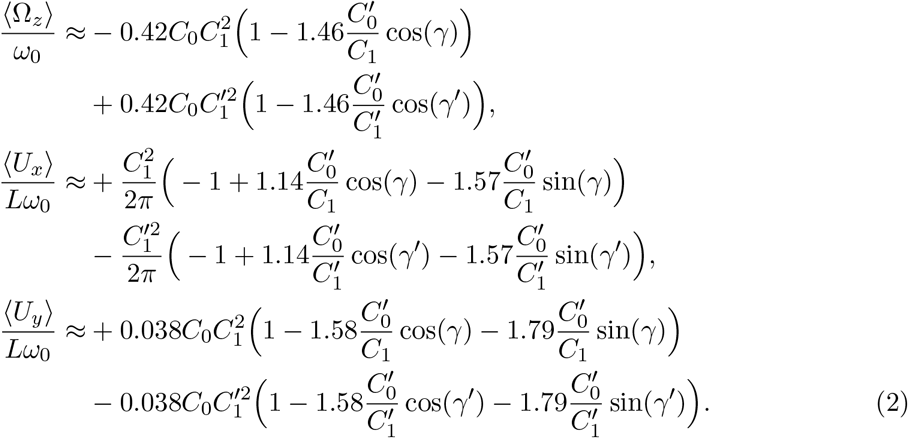

Here 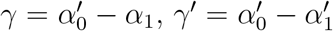, and *η* = *ζ*_∥_/*ζ*_⊥_ is assumed to be 0.5, where *ζ*_⊥_ and *ζ*_∥_ are transversal and parallel friction coefficients of a cylindrical segment of the axoneme (see Materials and Methods). The full form of expressions is shown in Eqs. S.1–S.3.

We comment on some properties of Eq. 2. For the axoneme to rotate, the presence of static curvature or the second harmonic (as shown in Eqs. S.6) is necessary to break the mirror symmetry, i.e. *C*_0_ ≠ 0 or *C*_2_ = 0 or both nonzero [27, 40]. If the wave pattern of the axoneme becomes its mirror image after half a period (i.e. *C*(*s, t*) = −*C*(*s, t* + *T*/2)), then the flagella will swim on average in a straight trajectory, which is also expected by symmetry. This is the case if only odd harmonics are kept in the wave form, i.e. *C*_1_ and 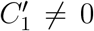. The reason is that the mean rotations achieved in the first and second half-periods, have the same magnitude but opposite signs and the net rotation sums up to zero. Equation 2 also shows that in the absence of even harmonics, 〈*U_y_*〉 = 0 and 〈*U_x_*〉 depends on the square of *C*_1_ and 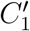 [51, 54, 55]. Furthermore, the term proportional to 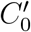 which describes global oscillations of the axonemal curvature, by itself can not contribute to the propulsion. However, once the time-symmetry is broken by other wave components, it contributes with a term proportional to 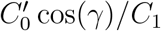 in Eq. 2 and can enhance the translational and rotational velocities with a factor of around 2 (assuming *γ* ~ *π* and 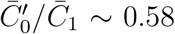 from Fig. 4B-C). Finally, the phase differences between first harmonics (*C*_1_ and 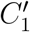) and the global oscillations (*γ* and *γ*′) also contribute to the translational and rotational velocities of the axoneme.

As mentioned above, the time-averaged mode amplitudes presented in Fig. 4B result in 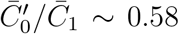. The corresponding phase values are shown in Fig. 4C, which gives 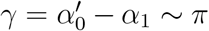 and 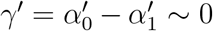. By substituting these numbers, Eq. 2 simplifies further to:

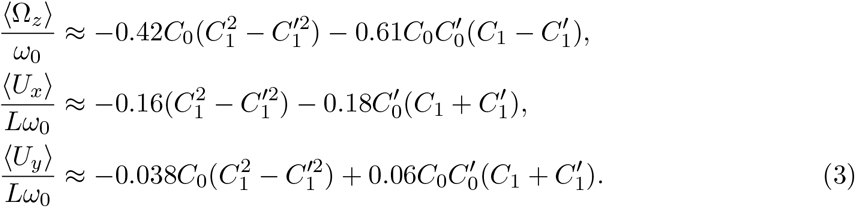

In parallel to our analytical approximations, we also performed numerical simulations using RFT with the simplified waveform presented in Eq. 1. We compared rotational and translational velocities obtained from our simulations with the analytical approximations presented in Eq. 2. Figure 6 shows a good agreement between the analytical approximations and the numerical simulations at small values of *C*_1_ and 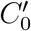, but deviations at larger values. These simulations are conducted at a fixed value of static curvature *C*_0_ = 0.04 and *C*_2_ = 0. Furthermore, an exemplary simulation in Fig. 7 highlights the effect of the global oscillations of the curvature 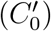 in increasing the translational and rotational velocity of a model axoneme, as also clearly visible in Fig. 6D (see Videos 5-6).

**Figure 6.**
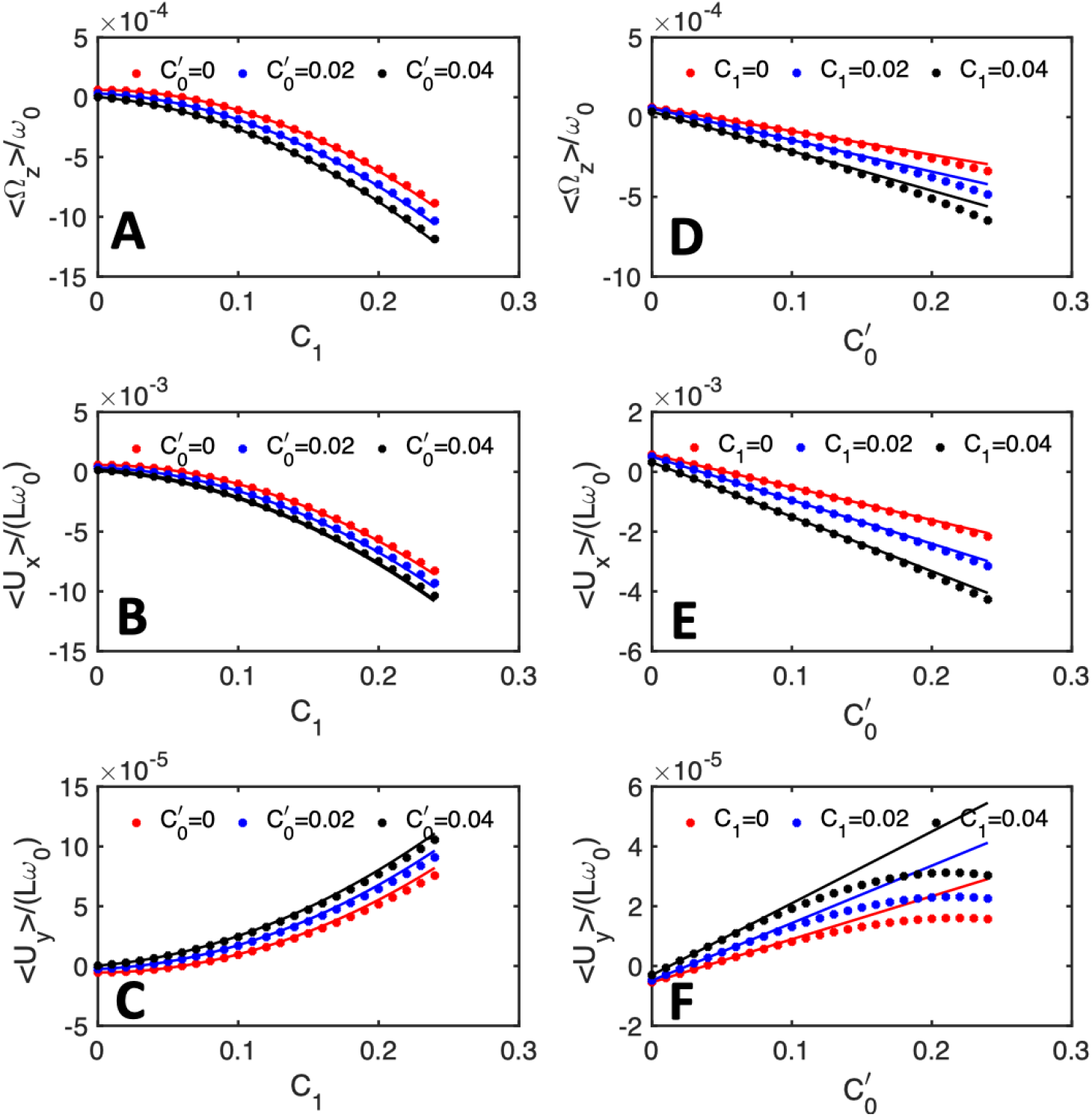
Rotational and translational velocities of a freely swimming axonemes as a function of *C*_1_ (A-C) and 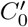 (D-F). Solid lines show the theoretical approximations as presented in Eq. 3 and the dots show the numerical simulations. As expected, there is a good agreement between analytical approximations and numerical simulations at small values of *C*_1_ and 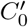, but deviations at larger values which point to the importance of nonlinear effects, not included in our analysis. The value of *C*_0_ (static curvature) is fixed at 0.04 in all the panels and *C*_2_ is zero.

**Figure 7.**
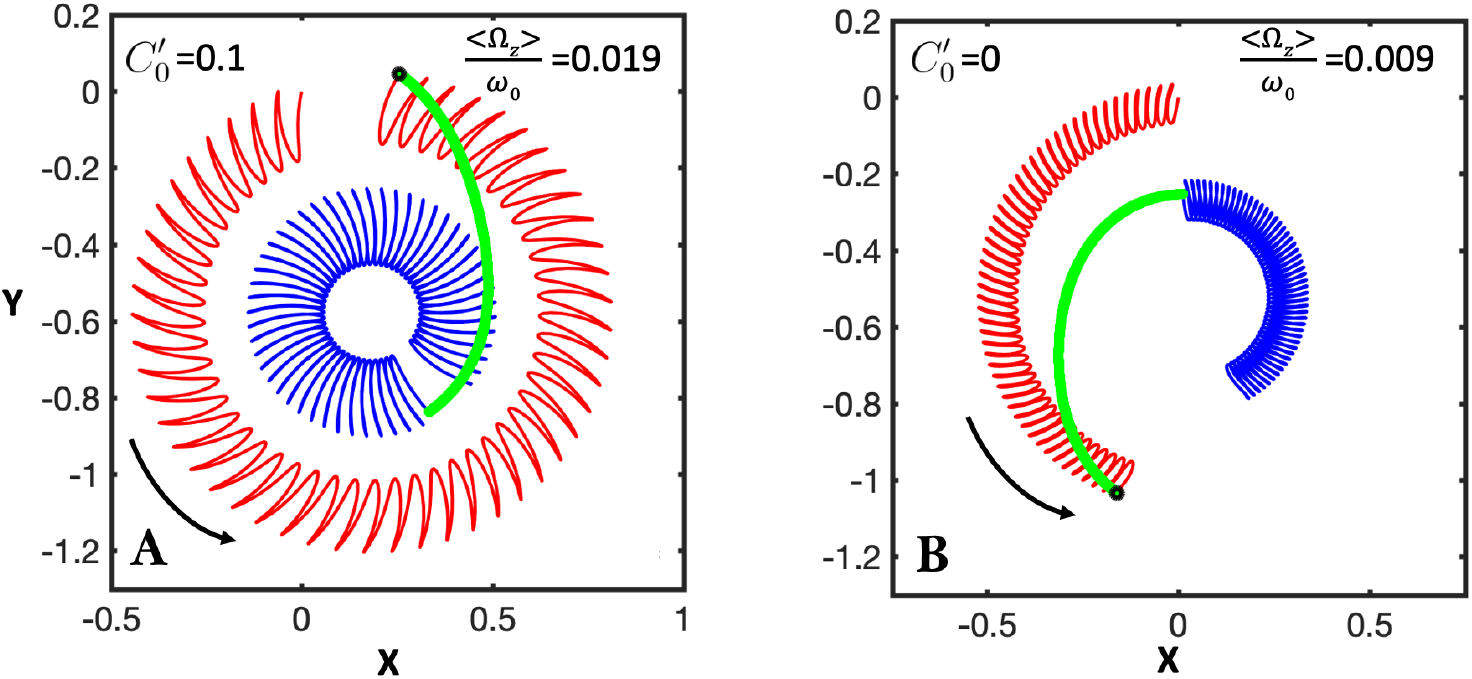
Simulations to show the effect of 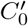 (amplitude of the global oscillations of the curvature) in increasing the rotational velocity. An axoneme with A) 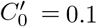 rotates roughly two times faster compared to another axoneme with 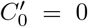. Simulations are done for the time interval of 998 msec. Red and blue trajectories correspond to the traces of basal and distal tips, respectively. Other parameters are: *f*_0_ = 50 Hz, *C*_0_ = −0.4, *C*_1_ = 0.2, *C*_2_ = 0, 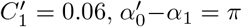, and 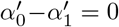. The values of translational and rotational velocities are measured as: A) (Ω_*z*_)/*ω*_0_=0.019, 〈*U_y_*〉/*Lω*_0_ = 0.0008, 〈*U_x_*〉/*Lω*_0_ = −0.015 and B) 〈Ω_*z*_〉/*ω*_0_ = 0.009, 〈*U_y_*〉/*Lω*_0_ = 0.0003, 〈*U_x_*〉/*Lω*_0_ = −0.008 (see also Videos 5-6).

### 2.3. Calcium reduces the intrinsic curvature of axonemes

To investigate the effect of calcium ions on the flagellar waveform, we performed experiments at different calcium concentrations and systematically measured the amplitude of the dominant contributing modes. As shown in Fig. 8, we changed the calcium concentration from 10^−4^ mM to 1 mM and quantified the curvature waves of the reactivated axonemes. Our mode decomposition analysis confirm that the static component of the curvature waves *C*_0_ shows a significant drop (~ 85%), as we increase [Ca^2+^] from 10^−4^ mM to 1 mM. This sudden decrease in *C*_0_ triggers a transition from circular swimming trajectory to a straight sperm-like path (see Fig. 8A and Video 7). This switch occurs at [Ca^2+^] around 0.02 mM. The wave amplitude C1 also shows a reduction of less than 50% as we increased the calcium concentration, but the remaining dominant modes 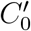 and *C*_2_ did not show a significant dependency on [Ca^2+^] (see Fig. 8C-D). Furthermore, we observed that the beat frequency of axonemes drops slightly at high calcium concentrations around 1 mM (Fig. 8E).

**Figure 8.**
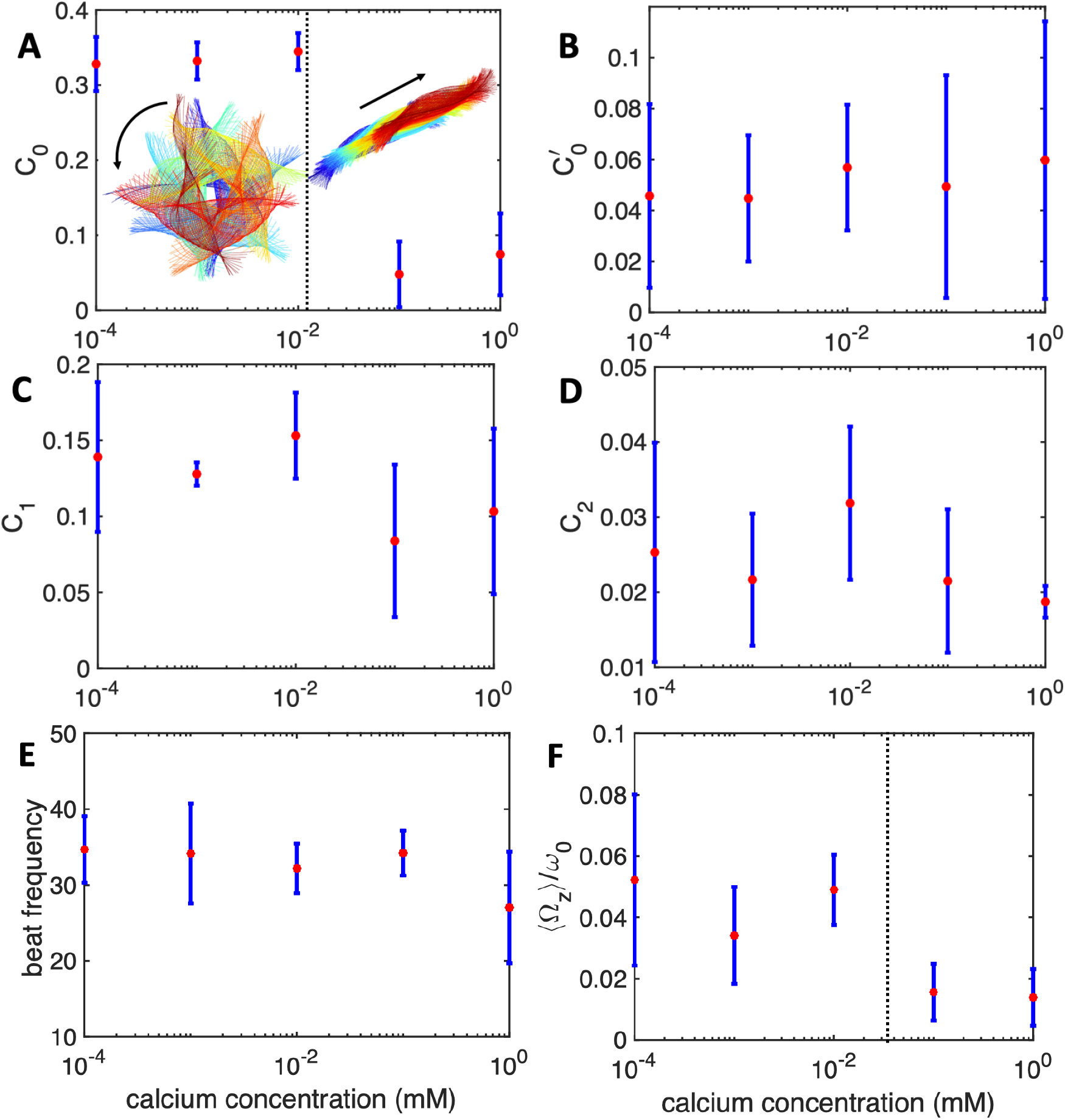
Experiments with calcium-supplemented reactivation buffer at different calcium concentrations. A) The static curvature *C*_0_ show a sudden drop from 0.3 to 0.05 at [Ca^2+^] around 0.02 mM which induces a switch from circular to straight swimming path. B) The mode amplitude describing the global oscillations of the axoneme 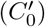 do not show a systematic trend as [Ca^2+^] has increased. C) The main wave amplitude *C*_1_ show a decrease with [Ca^2+^], but D) the second harmonic remains relatively unchanged. The error bars are calculated for five axonemes at each [Ca^2+^]. E) Beat frequency of axonemes drops slightly at [Ca^2+^] around 1 mM. F) Mean rotational velocity of axonemes decreases at [Ca^2+^] above 0.01 mM.

Similar to Fig. 5, we used *C*_0_, 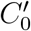, *C*_1_, 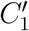 and *C*_2_ modes to reconstruct axonemal shapes for an exemplary axoneme at calcium concentration of 0.1 mM. As shown in Fig. 9, while removing *C*_2_ mode has almost no significant effect on fitting error (compare panels E-F with G-H), removing both 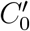 and 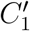 modes, increases the error around two times (compare panels F and J).

**Figure 9.**
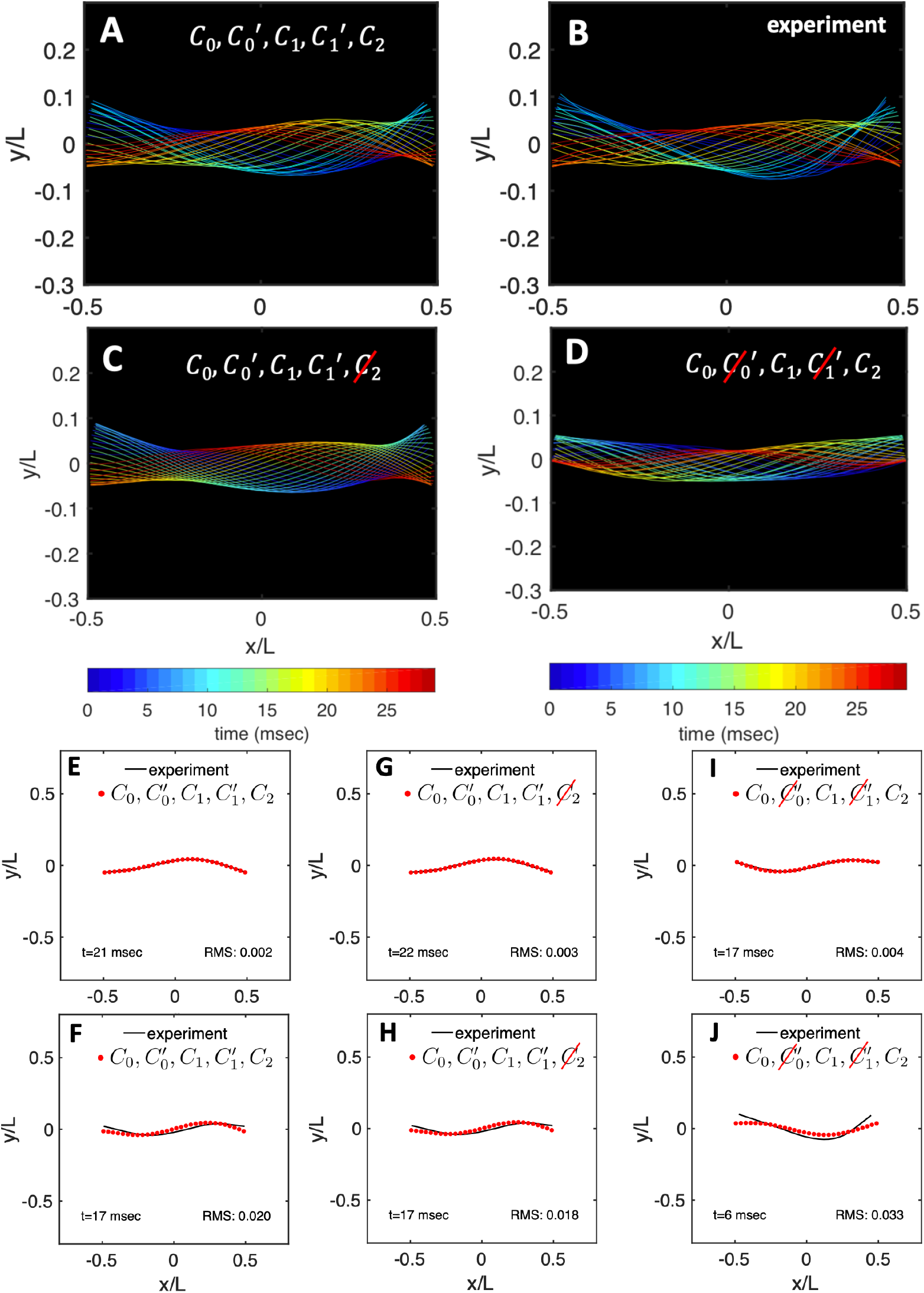
A) Five dominant modes, *C*_0_, 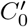, *C*_1_, 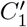 and *C*_2_, are used to reconstruct the experimental shapes shown in panel B. C) The shape reconstruction without the second harmonic *C*_2_ and D) without both back propagating wave 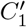 and global oscillations mode 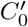. E-F) Exemplary fits of the best and the worst shape reconstructions shown in panel A with the root mean square (RMS) error of 0.002 and 0.020, respectively. G-F) The best and the worst shape reconstructions without *C*_2_ mode and I-J) without 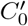 and 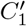 modes are shown. Calcium concentration is 0.1 mM; see Videos 8-10.

For reactivated axonemes in the presence of calcium, we also measured the mean rotational velocities. As shown in Fig. 8F, 〈Ω_*z*_)/*ω*_0_ reduces by increasing the calcium concentration, consistent with the observation that axonemes switch from circular to straight swimming trajectories. Furthermore, the analytical approximations in Eq. 2 predicts a linear dependency of 〈Ω_*z*_〉 on 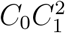 and 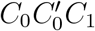. To examine the validity of this linear trend, we plot the mean rotational velocities of axonemes in dependence of 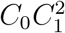 and 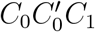 (see Figs. 10A-B). Consistent with our analytical prediction, a linear dependency was observed.

**Figure 10.**
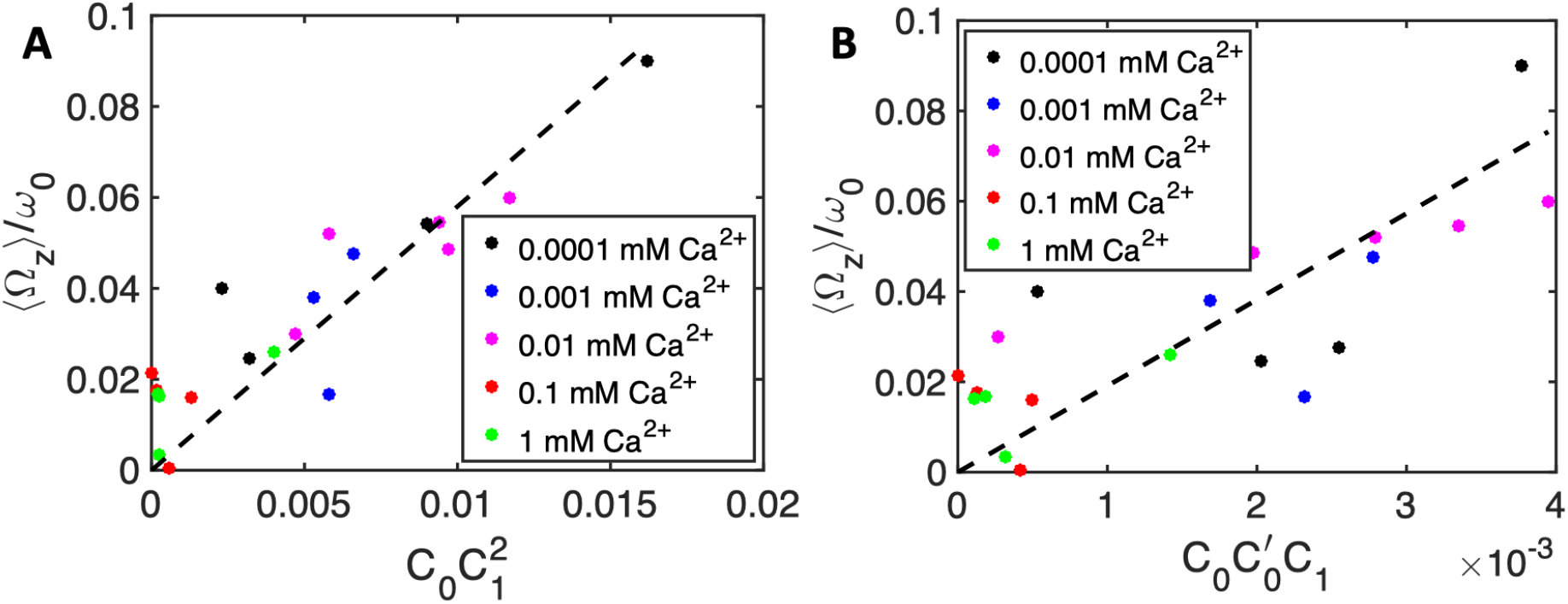
Mean rotational velocity of axonemes in dependence of A) 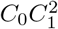 and B) 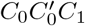. Black dashed lines represent a linear fit.

Finally, we note that in Figs. 10A-B, 〈Ω_*z*_〉 are calculated using the full experimental shapes based on RFT simulations with *η* = ζ_∥_/*ζ*_⊥_ = 0.5. The reason is that at high calcium concentrations with small rotational velocities of axonemes, Ω_*z*_(*t*) is very noisy and a direct measurement of 〈Ω_*z*_〉 is difficult. To check the validity of RFT in our system, we used our experimental data at zero or very low concentrations of calcium, where axonemes swim on circular paths and therefore a direct measurement of 〈Ω_*z*_〉 is possible. The comparison between the experimental measurements and RFT simulations is shown in Fig. 11A. The black line is a least square fit of slope 1, which provides a strong evidence for the validity of RFT in our experimental system. Furthermore, with regard to our mode analysis discussed previously, we mention that in our RFT simulations, the mean rotational velocity of axonemes depends on the number of modes which are considered for the shape reconstruction. Fig. 11B shows that 〈Ω_*z*_〉/*ω*_0_ obtained using RFT simulations, converges for *n* ≥ 15. The black dashed lines in Fig. 11B are the values of 〈Ω_*z*_〉^RFT^/*ω*_0_ obtained using full experimental shapes, i.e. infinite number of modes.

**Figure 11.**
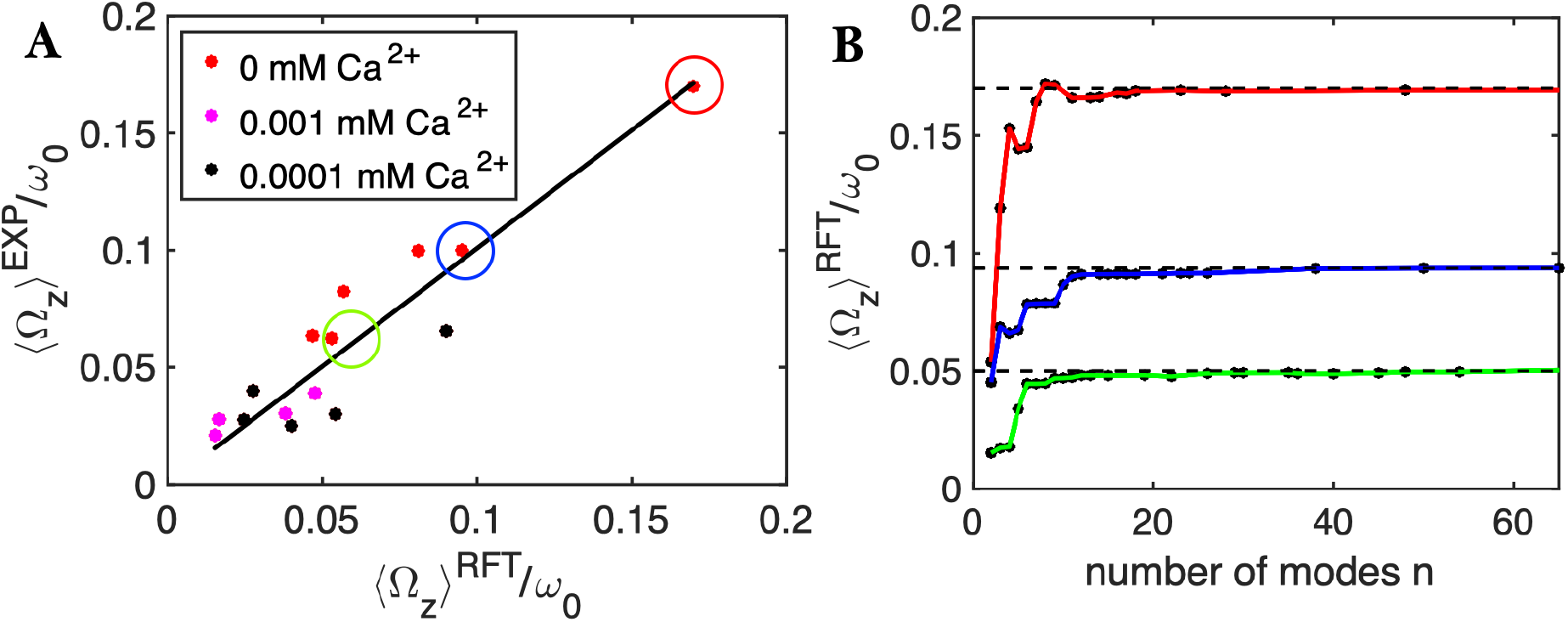
A) Comparison of experimental rotational velocities with corresponding values obtained by RFT simulations with full experimental shapes, i.e. infinite number of modes. The black line is a least square fit of slope 1. Each data point corresponds to one axoneme at a given calcium concentration. Circles show the data points which are used in panel B. B) The dependence of 〈Ω_*z*_〉^*RFT*^/*ω*_0_ on the number of modes considered for the shape reconstructions for the three axonemes, indicated by colored circles in panel A.

### 2.4. A pinned axoneme

In our experiments, attachment of axonemes to the substrate occurs either from the basal end or the distal tip due to the non-specific axoneme-substrate interactions (Fig. 12 and Video 11). In most cases, a pinned axoneme is free to rotate around the pinning point, so the total torque exerted on the axoneme is zero. Here, we consider an exemplary axoneme which is pinned from the basal end to the substrate (Fig. 12A-B). Curvature waves initiate at the proximal region and travel at the frequency of 18 Hz toward the distal tip (Fig. 12C). Similar to the free swimming axonemes, we measured an intrinsic curvature of ~ −0.19 *μ*m^−1^, as shown in Fig. 12D. The power spectrum of the “windowed” curvature shows dominant peaks at various modes, as highlighted in Fig. 13A. Notably, both the second harmonic (*C*_2_) and the mode describing the global oscillations 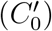 have similar amplitudes (Fig. 13B). Furthermore, phase information in Fig. 13C demonstrates that as in the case of the free axoneme in Fig. 4, modes 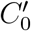 and 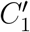 are almost in phase (compare 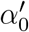 and 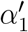), but *C*_1_ and 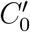 are almost out of phase (compare 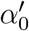 and *α*_1_).

**Figure 12.**
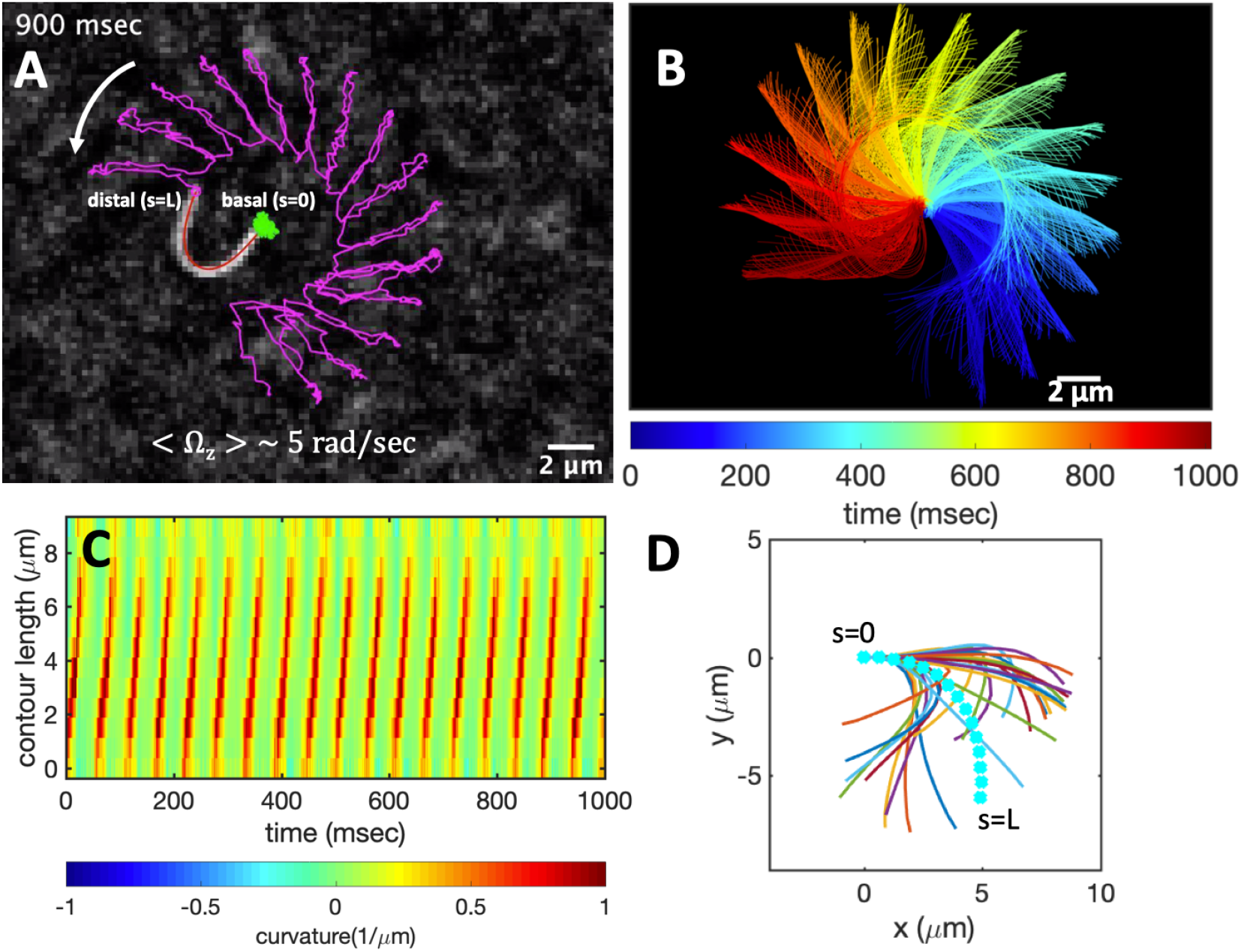
A-C) A pinned axoneme with beat frequency of 18 Hz rotates around the pinning point while the bending waves propagate from the basal toward the distal tip. D) The arc-shape mean shape shown in cyan color has mean curvature of −0.19 *μ*m^−1^. Concentration of ATP is 80 *μ*M; see also Video 11.

**Figure 13.**
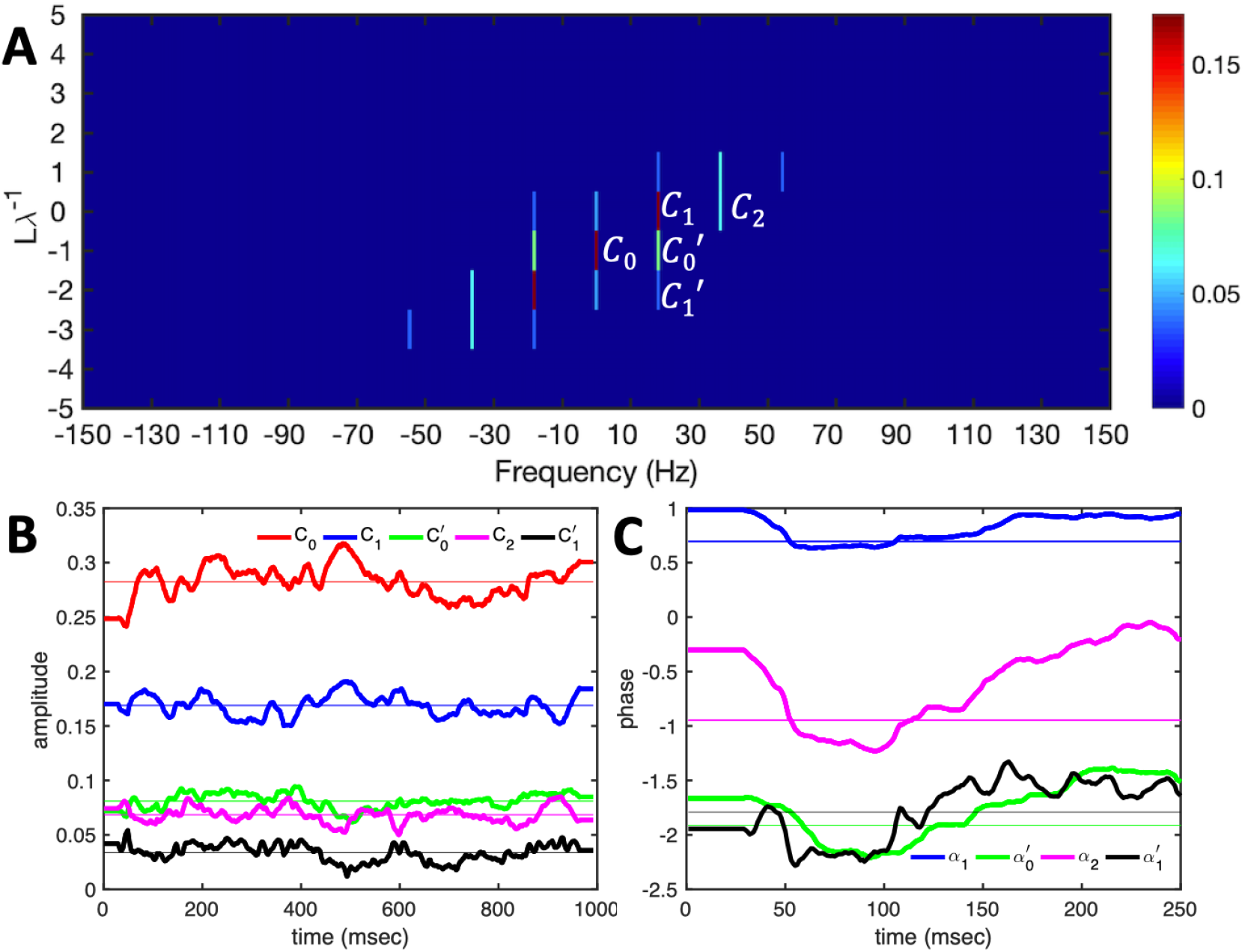
A) Power spectrum of the pinned axoneme presented in Fig. 12. Timevarying coefficients and the corresponding phase values are shown in panels B and C.

Following a similar procedure as for a free axoneme, we used the simplified waveform described in Eq. 1 and RFT to calculate the mean rotational velocity of a pinned axoneme to obtain (see Materials and Methods):

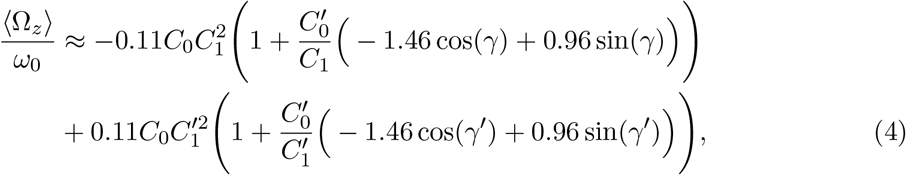

where with 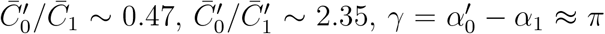 and 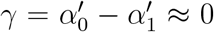 (see Fig. 13B-C), it further simplifies to:

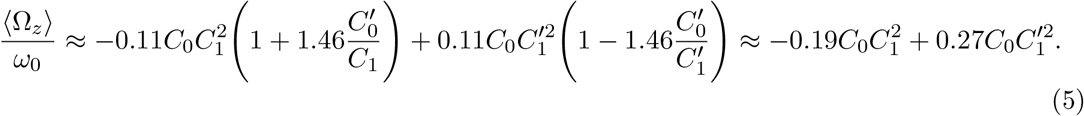

It is worth mentioning that since 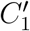 is almost five times smaller than *C*_1_, the first term has the dominant contribution in 〈Ω_*z*_〉. Note that the second harmonic also contributes in the rotational velocity of the axoneme (see supplemental information), but since the intrinsic curvature is around four times larger than the second harmonic, static curvature has the dominant contribution. Lastly, we emphasize that the term including the effect of global oscillations of the axoneme, contributes in enhancing the rotational velocity.

## 3. Discussion

In this work, we isolated flagella from green algae *C. reinhardtii* using dibucaine and removed the membrane by treatment of detached flagella with non-ionic detergents [8, 44]. The remaining 9+2 microtubule-based structure which lacks the membrane and basal bodies, can be reactivated by an ATP-supplemented buffer. Active axonemes show an asymmetric waveform predominantly in 2D which mostly resembles the forward swimming motion of flagella in intact cells [10, 12]. These asymmetrical beating patterns cause the axoneme to rotate stably around a position in the microscope’s field of view. We extracted the shape of axonemes by gradient vector flow technique [46, 47] and quantitatively described the flagellar beating patterns by dimensionless local curvature *C*(*s, t*) at time *t* and arc-length *s* along the axonemal length.

Power spectrum of the curvature waves demonstrates its most dominant peak at a static mode, highlighting the existence of an intrinsic curvature of axonemes around the value of ~ −0.2 *μ*m^−1^. To bend an axoneme to a circular arc, tangential axial forces generated by asymmetric distribution of dynein motors are required to induce a dynamic instability [30, 56–59]. Among different dynein motors, IDAb (inner dynein arm b) is possibly the only dynein which has an asymmetric distribution, both radially and longitudinally. Bui et.al. [60] have shown that IDAb is absent in the proximal region without being replaced by another dynein, and is predominantly localized at the distal tip with a radially asymmetric distribution. The depletion of IDAb from the proximal area might suggest its specific function in bending the axoneme. Future experiments with IDAb mutants are necessary to investigate the possible role of IDAb in inducing an intrinsic curvature in axonemes.

The second dominant peak of the power spectrum occurs at the fundamental beat frequency which describes a base-to-tip traveling wave component. However, our mode decomposition also highlights the coexistence of a tip-to-base propagating wave with an amplitude which is around five times reduced (compare the ratio between *C*_1_ and 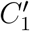 in Fig. 4B). These back-propagating waves also exist in the pinned axonemes. Simulations by Man et. al. [57] with a base-clamped model cilium have shown that by increasing axial stresses applied by the dynein motors, traveling waves which propagate from tip-to-base can switch direction. This switch depends on the boundary condition [57, 61, 62] and does not occur in the hinged filaments. These simulations are not consistent with our experiments, in which we observe the coexistence of propagating waves in both directions for free and hinged axonemes. It may be that the two opposing components of propagating waves in axonemes are generated by two sets of dynein motors exerting different magnitudes of forces at different locations along the axonemal length.

Another remarkable feature of the power spectrum is the coexistence of a time-symmetric mode, corresponding to the global oscillations of the axonemal curvature. The mechanisms underlying these global oscillations of the curvature is not known, but most probably involves oscillatory on-off activity of dynein molecular motors. These oscillations are reminiscent of a Hopf bifurcation at zero spatial frequency and is theoretically shown to occur at very high frequencies around 5-10 kHz in short filaments (~1-2 *μ*m in length) [30]. In our experiments, the global curvature oscillations have the same frequency as the fundamental beat frequency, and it coexists with other wave components. Future investigations are needed to understand the mechanisms by which such a time-symmetric motion is generated and tuned to enhance the flagellar propulsion.

Finally, in our experiments with calcium, we observed that among the constituting modes of the flagellar waveform, the static component is the most sensitive mode which reduces significantly (~10 times) at calcium concentrations above 0.02 mM. This reduction of static mode, triggers a switch from circular to straight swimming path, as previously observed in Refs. [31, 32]. High resolution structural information obtained by electron cryotomography [63], strongly supports the idea that calcium could regulate the transmission of mechanosignals. Gui et.al. [63] have identified a calcium responsive protein, called calmodulin, at the interface between RS1 (radial spoke 1) and IDAa (inner dynein arm a). It is suggested that calcium induces a conformational change in calmodulin, which can alter directly the wave pattern by affecting RS1-IDAa interaction. An alternative plausible mechanism is that calcium affects a calmodulin-like subunit (LC4) of the outer dynein arm (ODA) and consequently, influence the dynein behavior [64]. Further experiments are required to clarify the mechanism of dynein regulation and the precise role of calcium in shaping the flagellar waveform.

## 4. Materials and Methods

### 4.1. Isolation of axonemes from C. reinhardtii

We used wild-type *C. reinhardtii* cells, strain SAG 11-32b, to isolate flagella using dibucaine following the protocols in Refs. [44, 65]. First, we grew the cells axenically in TAP (tris-acetate-phosphate) medium on a 12h/12h day-night cycle. Cells release their flagella upon adding dibucaine, which we purify on a 25% sucrose cushion, and demembranate using detergent NP-40 in HMDEK solution (30 mM HEPES-KOH, 5 mM MgSO_4_, 1 mM DTT, 1 mM EGTA, 50 mM potassium acetate, pH 7.4) supplemented with 0.2 mM Pefabloc. The membrane-free axonemes were resuspended in HMDEK buffer plus 1% (w/v) polyethylene glycol (M_*w*_ = 20 kg mol^−1^), 0.2 mM Pefabloc and used freshly after isolation. To perform reactivation experiments, we diluted axonemes in HMDEKP reactivation buffer (HMDEK plus 1 % PEG) supplemented with 0.2 mM Pefabloc and 80 *μ*M ATP. Reactivation solution plus axonemes was infused into 100 *μ*m deep flow chambers, built from cleaned glass and double-sided tape. To avoid axoneme attachment to the substrate, glass slides are coated with 2 mg/mL casein solved in HMDEKP. For calcium experiments, HMDEKP reactivation buffer is supplemented with calcium at different concentrations.

### 4.2. Axoneme Tracking

We used high-frame phase-contrast microscopy to analyze fast beating dynamics of axonemes (100X objective, 1000 fps). First, we invert phase-contrast images and subtracted the mean-intensity of the time series to increase the signal to noise ratio [26]. Next, we applied a Gaussian filter to smooth the images. Tracking of axonemes is done using gradient vector flow (GVF) technique [46, 47, 50]. For the first frame, we select a region of interest that should contain only one actively beating axoneme (see Fig. S2). Then, we initialize the snake by drawing a line polygon along the contour of the axoneme in the first frame. This polygon is interpolated at *N* equally spaced points and used as a starting parameter for the snake. The GVF is calculated using the GVF regularization coefficient *μ* = 0.1 with 20 iterations. The snake is then deformed according to the GVF where we have adapted the original algorithm by Xu and Prince for open boundary conditions. We obtain positions of *N* points along the contour length *s* of the filament so that *s* = 0 is the basal end and *s* = *L* is the distal side, where *L* is the total contour length of filament. The position of filament at *s_i_* is denoted by *r*(*s_i_*) = (*x*(*s_i_*), *y*(*s_i_*)).

### 4.3. Resistive force theory and calculations of mean translational and rotational velocities of a free axoneme

Biological microorganisms swim with flagella and cilia in the world of “low Reynolds number” where they experience viscous forces many orders of magnitude larger than inertial forces [42, 48]. In this world where inertia is negligible, Newton’s law becomes an instantaneous balance between external and fluid forces and torques exerted on the swimmer, i.e. 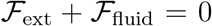 and 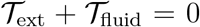. The force 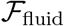 and torque 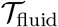 exerted by the fluid on the axoneme can be written as:

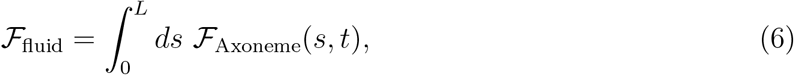

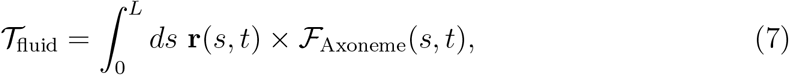

where the integrals over the contour length *L* of the axoneme calculate the total hydrodynamic force and torque exerted by the fluid on the axoneme. ATP-reactivated axonemes show oscillatory shape deformations. At any given time, we consider an axoneme as a solid body with unknown translational and rotational velocities **U**(*t*) and Ω(*t*), yet to be determined. 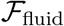 and 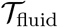 can be separated into propulsive part due to the relative shape deformation of the axoneme in body-fixed frame and drag part [66]:

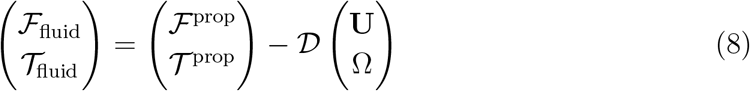

where 6×6 geometry-dependent drag matrix of the axoneme 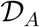 is symmetric and nonsingular (invertible). We also note that a freely swimming axoneme experiences no external forces and torques, thus 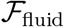 and 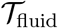 must vanish. Further, since swimming effectively occurs in 2D, 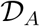 is reduced to a 3×3 matrix and Eq. 8 can be reformulated as:

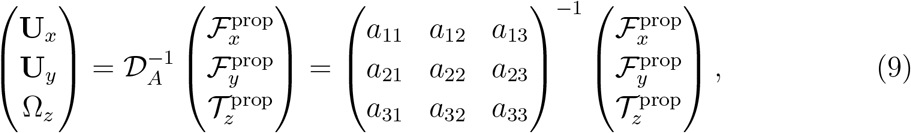

which we use to calculate translational and rotational velocities of the swimmer after determining the drag matrices 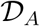 and the propulsive forces and torque 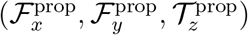.

We calculate 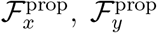 and 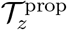 which are propulsive forces and torque due to the shape deformations of the axoneme in the body-fixed frame by selecting the basal end of the axoneme as the origin of the swimmer-fixed frame. As shown in Fig. 2A, we define the local tangent vector at contour length *s* = 0 as 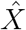 direction, the local normal vector 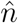 as 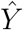 direction, and assume that 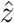 and 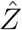 are parallel. Let us define *θ*_0_(*t*) = *θ*(*s* = 0, *t*) as the angle between 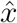 and 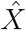 which gives the velocity of the basal end in the laboratory frame as 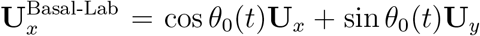 and 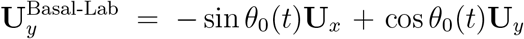. Furthermore, note that the instantaneous velocity of the axoneme in the lab frame is given by **u** = **U** + Ω × **r**(*s, t*) + **u**′, where **u**′ is the deformation velocity of flagella in the body-fixed frame, *U* = (**U**_*x*_, **U**_*y*_, 0) and Ω = (0, 0, Ω_*z*_) with Ω_*z*_ = *dθ*_0_(*t*)/*dt*.

To calculate 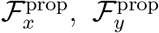 and 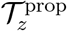 for a given beating pattern of axoneme in the body-fixed frame, we used classical framework of resistive force theory (RFT) which neglects long-range hydrodynamic interactions between different parts of the axoneme [52, 53]. In this theory, axoneme is divided into small cylindrical segments moving with velocity **u**′(*s, t*) in the body-frame and propulsive force 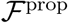 is proportional to the local centerline velocity components of each segment in parallel and perpendicular directions:

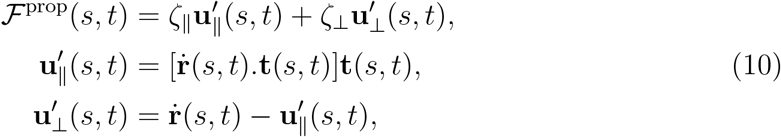

where 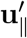 and 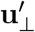 are the projection of the local velocity on the directions parallel and perpendicular to the axoneme. The friction coefficients in parallel and perpendicular directions are defined as *ζ*_∥_ = 4*πμ*/(ln(2*L*/*a*) + 0.5) and *ζ*_⊥_ = 2*ζ*_∥_, respectively. This anisotropy indicates that to obtain the same velocity, one would need to apply a force in the perpendicular direction twice as large as that in the parallel direction [52, 53]. Axoneme is a filament of length ~ 10 *μ*m and radius *a* ~ 100 nm. For a water-like environment with viscosity *μ* = 0.96 pN msec/*μ*m^2^, we obtain *ζ*_∥_ ~ 2.1 pN msec/*μ*m^2^.

### 4.4. Approximation of the mean rotational velocity of a pinned axoneme

Consider an axoneme which is attached from the basal end to the substrate and rotates around the pinning point (see Fig. 12A). We select the pinning point of the axoneme as the origin of the swimmer-fixed frame, and define tangent and normal vectors at *s* = 0 as the coordinate system in the swimmer-fixed frame. Since the axoneme is free to rotate but not to translate, the total torque 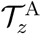 exerted on the axoneme is zero but the total force 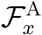 and 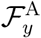 is non-zero. Reformulating in terms of Eq. 8, we have:

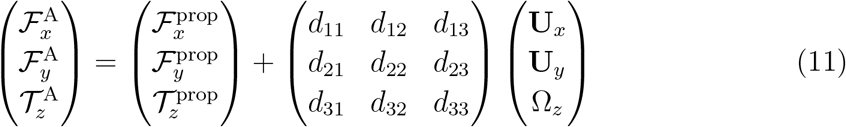

which by imposing the constrains of **U**_*x*_ = 0, **U**_*y*_ = 0 and 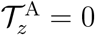 simplifies to:

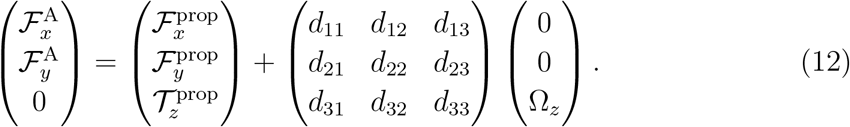

Using this equation, we can calculate the instantaneous rotational velocity of a pinned axoneme as:

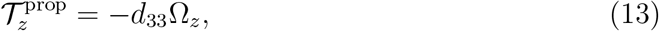

where

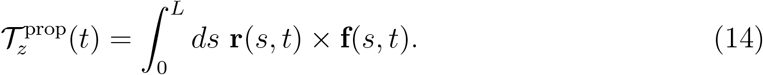

Here **f**(*s, t*) is calculated using Eq. 10 based on the simplified waveform defined in Eq. 1. To calculate mean rotational velocity for a pinned axoneme, we average over one beating cycle:

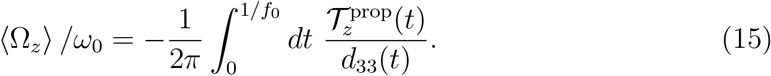

## Acknowledgment

The authors acknowledge V. Geyer, G. Gompper, J. Elgeti, and J. Molacek, and F. Nordsiek for valuable comments, S. Goli Pozveh and Y. Su for their support and S. Romanowsky, M. Muller and K. Gunkel for technical assistance. A.G. and A.B. acknowledge MaxSynBio Consortium, which is jointly funded by the Federal Ministry of Education and Research of Germany and the Max Planck Society. R.A. acknowledge support from the European Union’s Horizon 2020 research and innovation programme under grant agreement MAMI No. 766007. The authors also thank M. Lorenz and S. Bank at the Göttingen Algae Culture Collection (SAG) for providing the *C. reinhardtii* wild type strain SAG 11-32b.

## 5. Supplementary Materials

### 5.1. Full expressions of the mean rotational and translational velocities of a free and a pinned axoneme

For a freely swimming axoneme, the mean rotational velocity looks as following:

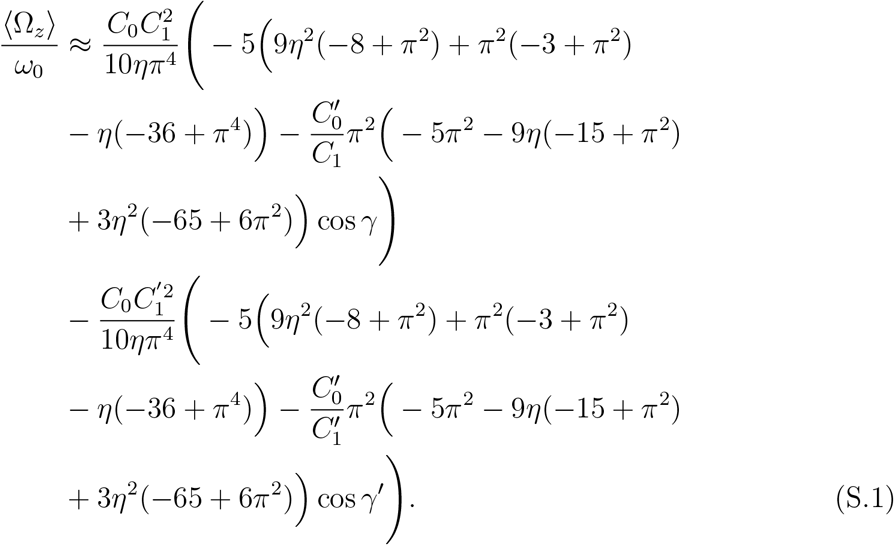

Similarly, we obtain:

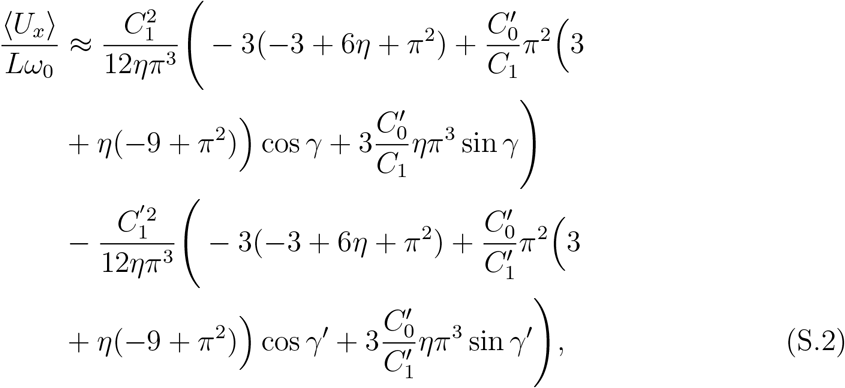

and

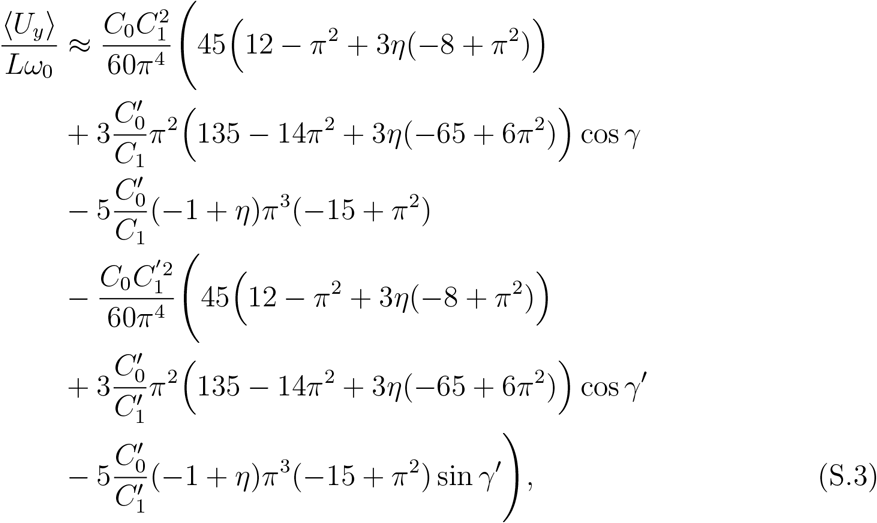

where with *η* = *ζ*_∥_/*ζ*_⊥_ = 0.5 [52, 53], they simplify to Eq. 2. Here *ζ*_⊥_ and *ζ*_∥_ are transversal and parallel friction coefficients of a cylindrical segment of the axoneme. Here 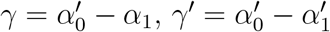.

For a pinned filament, before performing the integration in Eq. 15 over time, we expand the ratio of 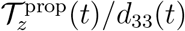 up to the first order in *C*_0_ = *κ*_0_*L*/(2*π*) and 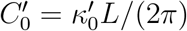 and second order in *C*_1_ = *κ*_1_*L*/(2*π*) and 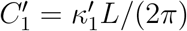, to obtain:

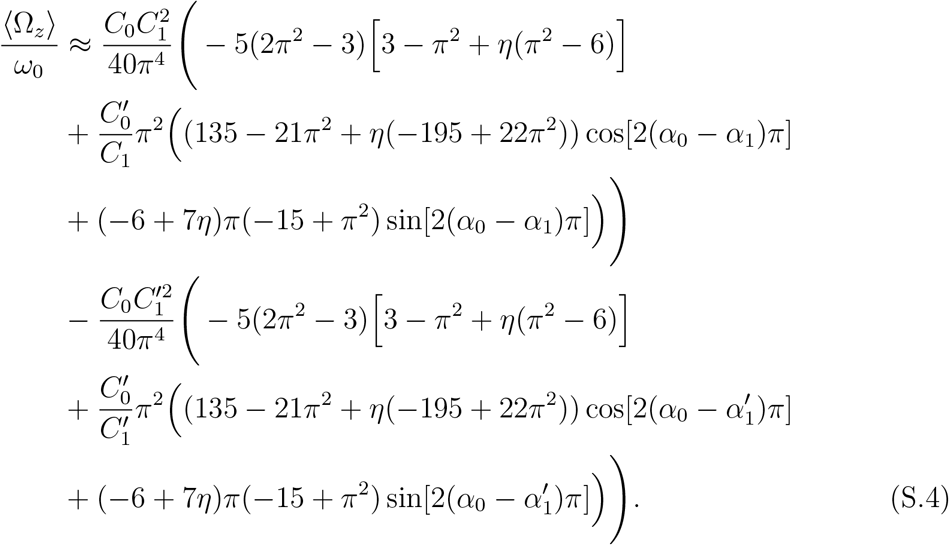

Assuming *η* = *ζ*_∥_/*ζ*_⊥_ = 0.5 [52, 53], Eq. S.4 simplifies to Eq. 4.

### 5.2. Effect of the second harmonic

Let us assume the following wave form which also includes the effect of the second harmonic:

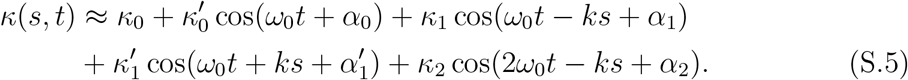

Following the same procedure as described in Materials and Methods, Sec. 4.3, we obtain:

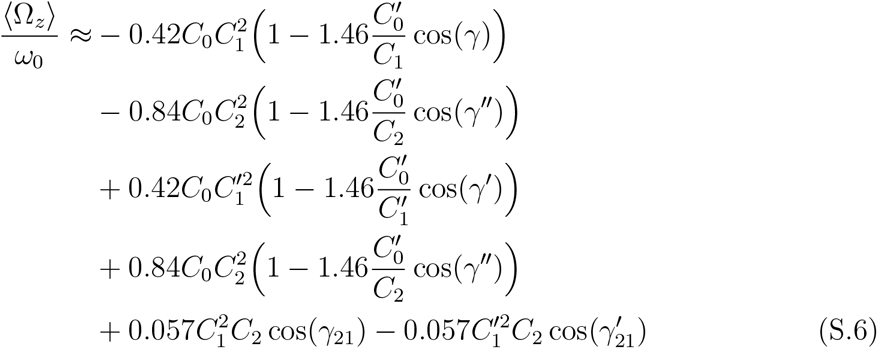

where 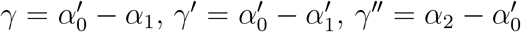, *γ*_21_ = *α*_2_ – *α*_0_ and 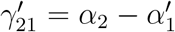. Similarly, for translational velocities, we get:

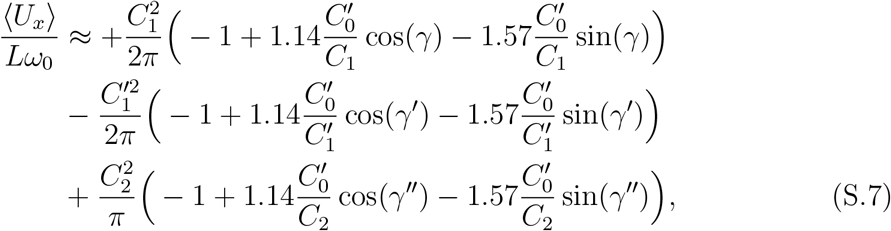

and

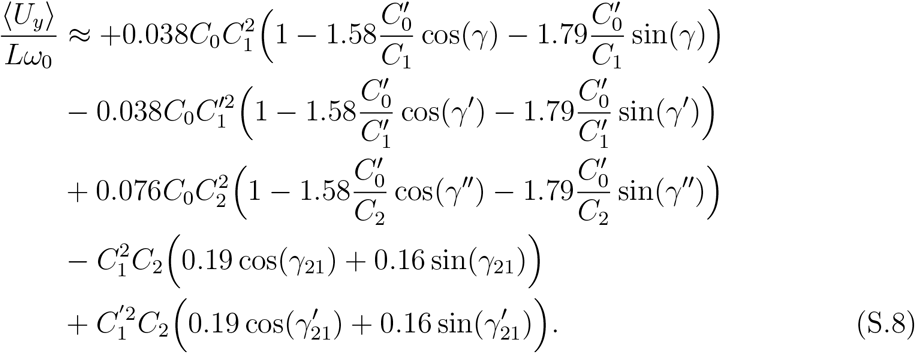

**Figure S1.**
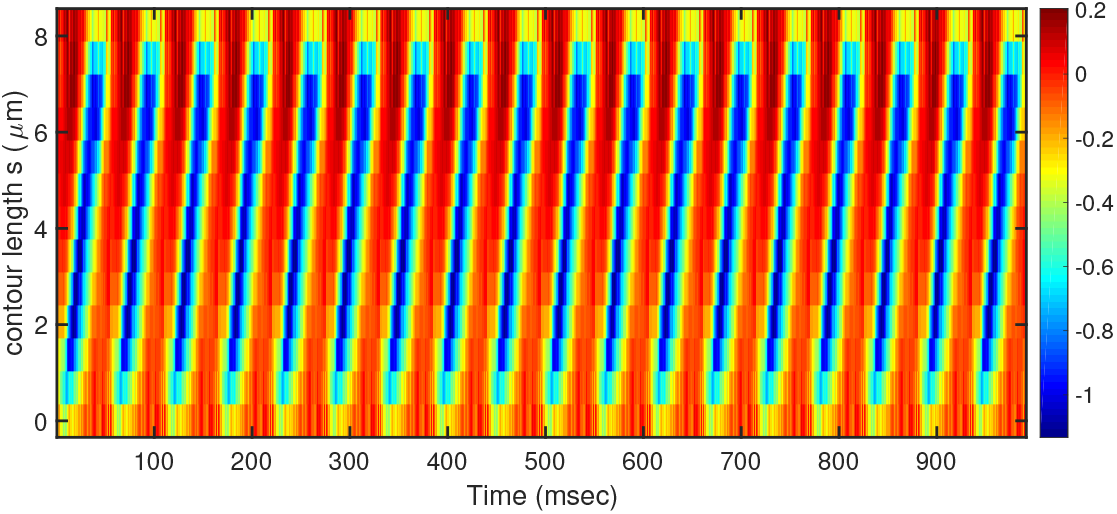
Windowed curvature of the data presented in Fig. 2C. Last beat cycle is repeated integer number (*n* = 17) of times.

**Figure S2.**
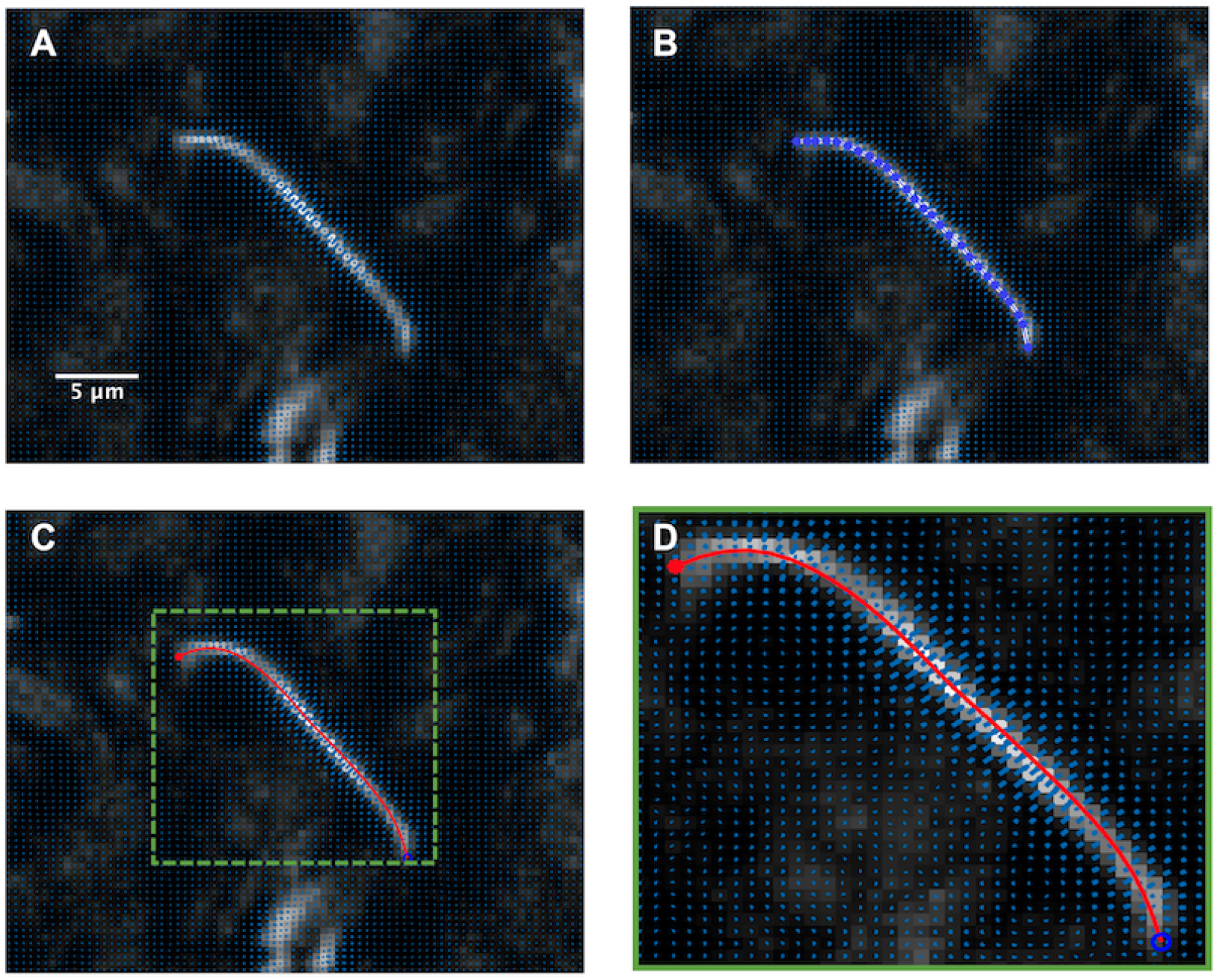
A) Gradient vector flow calculated at the vicinity of an exemplary axoneme. B) The initial selection of a polygon for the first frame which deforms according to the gradient vector flow shown in blue arrows. C) The final tracked shape of an axoneme. D) A zoomed-in image of the area shown by a green dashed box in panel C.

